# Single-cell analysis of 2,3,7,8-tetrachlorodibenzo-*p*-dioxin (TCDD)-treated murine skin demonstrates that sebaceous gland differentiation precedes *Ahr*-dependent seboatrophy with elevated expression of infundibular *Blimp1*

**DOI:** 10.64898/2026.07.22.740060

**Authors:** Syeda Mashnia Tasnim, Subhash Solanki, Jyoti Bhuju, Lauren Thompson, Omar Skalli, Elizabeth A. Grice, Carrie Hayes Sutter, Thomas R. Sutter

## Abstract

In humans, 2,3,7,8-tetrachlorodibenzo-*p*-dioxin (TCDD) induces chloracne, a skin condition that presents with acanthosis, hyperkeratosis, comedones, and sebaceous gland (SG) atrophy (seboatrophy). Although chloracne-like phenotypes have been reported in TCDD-treated mice, the underlying mechanisms remain poorly understood. Previous studies showed that TCDD-induced CYP1A1 protein is expressed in LRIG1+ progenitor cells in hair follicles, suggesting that TCDD targets specific cell populations within the pilosebaceous unit. To explore the effects of TCDD on the epidermis and pilosebaceous unit, we analyzed single-cell RNA expression in wild-type and *Ahr*-null mice at postnatal day 21 (P21) following *in utero* and lactational exposure. The results showed that TCDD preferentially induced the AHR target genes *Cyp1a1* and *Cyp1b1* in the lower infundibulum and subjacent junctional zone overlapping the LRIG1+ progenitor cell niche. TCDD also caused *Ahr*-dependent seboatrophy, accompanied by increased expression of *Blimp1*, a transcriptional repressor that regulates SG size. A second site of *Cyp1a1* induction was the SG, where *Cyp1a1* was markedly elevated in the basal proliferating cells and immature sebocytes. In a 3-day topical exposure study of early effects, TCDD produced a dose-dependent increase of *Cyp1a1* expression in the SG that included the more differentiated sebocytes. This response was accompanied by expansion of the *Scd1*-positive area, elevated Nile Red lipid staining, and an increased number of *Blimp1*-high sebocytes, demonstrating that TCDD enhanced SG differentiation and lipid production *in vivo*. These changes preceded the onset of *Ahr*-dependent seboatrophy, providing new insight into the cellular and molecular events underlying chloracne pathogenesis.

## Introduction

The skin is part of the integumentary system and the largest organ of the body with an area of about 2m^2^ (Wysocki 1999). It is an essential barrier organ that protects against the external environment, acts as a sensory organ, prevents fluid loss, and helps to maintain body temperature (Hwa et al. 2011). The pilosebaceous unit is a dynamic component of the skin that includes both the hair follicle and its attached sebaceous gland (s) (SG) and is contiguous with the interfollicular epidermis (IE). SGs are unique acinar gland adnexa of hair follicles that are comprised of both epithelial cells and lipid filled sebocytes that secrete an oily and waxy substance called sebum (Schneider and Paus 2010). Sebum provides the skin with moisture, a protective barrier and thermoregulation (Zouboulis et al. 2008). Proliferating basal epithelial cells form the outer layer of the SG and the transitional basal cells in the peripheral zone give rise to the undifferentiated sebocytes. As sebocytes move, towards the center of the gland and outward, they become more differentiated and produce and release sebum into the sebaceous duct via holocrine secretion (Niemann 2009; Niemann and Horsley 2012). In mice, LRIG1+ progenitor cells in the junctional zone (JZ) of the upper portion of the hair follicle give rise to the SG and its sebocytes during hair follicle morphogenesis and differentiation (Frances and Niemann 2012).

The aryl hydrocarbon receptor (AHR) is a ligand-activated transcription factor that regulates skin homeostasis, xenobiotic metabolism, and barrier integrity (Kennedy et al. 2013; Muenyi et al. 2014; Sutter et al. 2011). 2,3,7,8-Tetrachlorodibenzo-*p*-dioxin (TCDD) is an environmental pollutant and a high affinity ligand of the AHR, exerting toxic effects through the receptor. Deleterious effects of TCDD include reproductive toxicity, immune suppression, endocrine disruption and cancer (Birnbaum 1994). Chloracne, a skin disorder characterized by acanthosis, hyperkeratosis, skin lesions, comedones and metaplastic changes to sebaceous glands is an outcome of TCDD-mediated toxicity in humans (Knutson and Poland 1982; Panteleyev and Bickers 2006; Suskind 1985). The poisoning of Viktor Yushchenko with TCDD is the most extensively studied case of chloracne in recent years. In this case, sebaceous gland atrophy was accompanied by the formation of cysts that expressed CYP1A1 in the walls of these lesions identified as hamartomas (Saurat et al. 2012). While chloracne was initially thought to be restricted to the sebaceous and meibomian glands of primates, several *in vivo* studies demonstrate sebaceous gland atrophy in both hairless and haired mice (Bhuju et al. 2021; Fontao et al. 2018; Muku et al. 2019; Puhvel et al. 1982).

The expression of certain cytochrome P450 genes such as *Cyp1a1* and *Cyp1b1* serve as biomarkers of AHR activation. In mice treated with TCDD, CYP1A1 protein has a very different expression pattern in comparison to the AHR protein. Whereas the expression of the AHR occurs throughout the pilosebaceous unit, CYP1A1 induction colocalizes to the follicular region of the lower infundibulum and junctional zone, where the LRIG1+ SG progenitor cells reside (Bhuju et al. 2021; Fontao et al. 2018). This unique colocalization of CYP1A1 and LRIG1 proteins led the authors to propose that TCDD may selectively target this specific stem cell population to affect seboatrophy (Bhuju et al. 2021; Fontao et al. 2018).

The development of the SG depends, in part, on MYC. MYC plays a role in the cell fate decision and proliferation of the LRIG1+ cells to form the SG (Honeycutt and Roop 2004). MYC also plays a critical role in the differentiation of the SG transitional basal cells into sebocytes (Cottle et al. 2013; Honeycutt and Roop 2004). Overexpression of MYC induces sebaceous hyperplasia (Arnold and Watt 2001), whereas inactivation of *Myc* causes impaired secretion of sebum (Zanet et al. 2005). As such, MYC contributes to both SG gland morphogenesis and cell differentiation. BLIMP1, a repressor of MYC, is reported to be expressed in the terminally differentiated cells of epidermis, hair follicle and SG (Kretzschmar et al. 2014). Loss of *Blimp1* results in overexpression of MYC coupled with an increased number of sebocytes and SG enlargement (Horsley et al. 2006). This suggests that the interplay of MYC and BLIMP1 regulates SG homeostasis.

Considering the colocalization of AHR biomarkers with LRIG1+ SG progenitor cells, and the interactions between LRIG1, MYC and BLIMP1 for maintaining SG homeostasis, we investigated whether TCDD activates the AHR in a cell-specific context to alter sebaceous gland morphology and differentiation. Here, we utilized a quantitative single-cell *in situ* hybridization approach to further elucidate the regional and cell-selective targets of TCDD, and to better understand the effects of TCDD on the sebaceous gland that may contribute to the progression of chloracne.

## Materials and Methods

### Mice

Mice were obtained from Jackson Laboratory (Bar Harbor, ME, USA). C57BL/6J (strain# 000664) *Ahr* +/+ mice were used as wild-type mice or bred with the *Ahr*-null strain, B6.129-Ahr*^tm1Bra^*/J mice (Schmidt et al. 1996) (strain# 002831). Animals were housed in clean transparent plastic cages and maintained on a 12:12-hour light-dark cycle, temperature (18-26 °C), and humidity (40-60%) in a controlled room of the animal care facility. Mice were acclimated to Teklad Global 16% Protein Rodent Diet 2016 (Harlan Teklad, Madison, WI, USA) at least one week before experiments were initiated. Breeding and treatment of the mice followed previously published protocols (Bhuju et al. 2021; Muenyi et al. 2014). Heterozygous *Ahr* mice were bred, and the *Ahr* genotypes were determined by PCR and confirmed by immunohistochemical analysis **(Fig. S1)**. For the post-natal day 21 (P21) study, heterozygous populations from different litters were used for crossing to avoid bias. Pregnant, time-mated dams were treated with TCDD, as described below and homozygote (+/+ or -/-) pups were recruited into experiments. The breeding and treatment of the mice followed previously published methods (Bhuju et al. 2021; Muenyi et al. 2014). All animal research protocols were reviewed and approved by the Institutional Animal Care and Use Committee (IACUC) of the University of Memphis.

### Treatments

#### i) P21 study

Treatments with TCDD were performed as previously described (Bhuju et al. 2021). On E12, pregnant dams were weighed and randomly assigned to groups to receive either corn oil (control vehicle) or 5 µg/kg body weight TCDD in 100 µL corn oil via a single oral gavage (Bhuju et al. 2021; Muenyi et al. 2014). Four experimental groups were produced (n = 6-8) based on treatment and genotype: control wild-type +/+, treated wild-type +/+, control *Ahr*-null -/-, and treated *Ahr*-null (-/-). Pups remained with their mothers until postnatal day 21 (P21), when they were euthanized for tissue collection.

#### ii) 3-day study

Eight-week-old C57BL/6J mice were used in this study. Mice were divided into 4 groups (n = 8/group, 4 males and 4 females) and topically treated with control vehicle [DMSO:acetone (1:9)] (D2650, Sigma-Aldrich, Milwaukee, WI, USA) or TCDD (ED-901-B, Cambridge Isotope Laboratories, Tewksbury, MA, USA), at concentrations of [0.00003% (300 ng/mL), 0.000003% (30 ng/mL), or 0.0000003% (3 ng/mL) in vehicle]. The doses were selected based on our previous study (Solanki et al. 2026). Mice were topically treated (100µL on the shaved dorsal back) once per day for 3 days and the skin was collected on the day after the last treatment.

### Tissue harvesting

After euthanasia, skin tissue was collected and fixed in 10% neutral buffered formalin. The next day, tissues were transferred to 70% ethanol at 4°C overnight and then processed and embedded in paraffin. For frozen sections, a second set of tissues was transferred to 20% (w/v) sucrose at 4°C overnight and embedded in OCT compound for cryostat sectioning.

### Immunohistochemistry

Immunoperoxidase staining was performed on 5 µm formalin-fixed paraffin-embedded (FFPE) skin tissue sections using a biotin/streptavidin detection system (VECTASTAIN® Universal Quick HRP Kit, PK-8800, Vector Laboratories, Burlingame, CA, USA) with DAB (DAB substrate kit, SK-4100, Vector Laboratories, Burlingame, CA, USA). Following deparaffinization, the sections were placed in 0.3% hydrogen peroxide in methanol for 30 minutes to block the endogenous peroxidase activity. Next, the tissue sections were incubated in 10% normal horse serum (NHS) blocking solution for 30 minutes and then dipped in 1X PBS. Hydrophobic barriers were drawn around each tissue section with a pen. The slides were placed in a humid chamber and incubated at 4°C with the primary antibody: anti-AHR (1:200, 28727-1-AP, Proteintech, Rosemont, IL, USA). After the incubation period, the slides were washed in TBS-T and PBS, each for 4 minutes. Next, the slides were incubated with biotinylated universal secondary antibody (1:20 dilution in PBS/NHS), followed by incubation with streptavidin-peroxidase (at a 1:45 dilution in PBS) for 5 minutes. Between each incubation, slides were washed in TBS-T and PBS, each for 4 minutes. Signal detection was performed by incubating the sections for 35 seconds with the chromogenic DAB peroxidase substrate. Following two washes of 1 minute each in tap water, the slides were counterstained with Mayer’s hematoxylin (26043-06, Electron Microscopy Sciences, Morgantown, PA, USA) for 30 seconds, rinsed with tap water and dipped 10 times in an acid rinse solution (2 mL glacial acetic acid in 98 mL DI water), 10 times in tap water, 1 minute in a bluing solution (30% stock NH4OH diluted in 70% ethanol), and 10 dips in tap water. Finally, the slides were dehydrated in different concentrations of ethanol, cleared in xylene, and then cover-slipped using a xylene-based mounting medium. Images were acquired using Olympus BX63 bright field microscope.

### Oil Red O staining

Lipid accumulation was assessed on 10 µm frozen tissue sections stained with Oil Red O (26503-02, Electron Microscopy Science, Morgantown, PA, USA). A working solution was prepared by mixing Oil Red O stock (0.5% in isopropanol) with DI water in a 3:2 (v/v) ratio. The frozen sections were rinsed in 60% isopropanol for 2-3 minutes and stained in the Oil Red O working solution for 10 minutes. After three rinses with 60% isopropanol (1 minute each), sections were counterstained with Mayer’s hematoxylin for 3 seconds. Slides were then washed in DI water three times, 2 minutes each, and cover-slipped using a water based mounting medium. Images were acquired using Olympus BX63 bright field microscope.

### Nile Red staining

SG lipid content was assessed on 10 µm frozen tissue sections by staining with the lipophilic dye Nile Red (N3013, Sigma-Aldrich), diluted to a 300 nM working solution in PBS. Tissue sections were incubated with Nile Red for 15 minutes at room temperature, protected from light. The slides were then rinsed once in PBS, followed by two 5 minutes washes in PBS. A coverslip was placed onto the tissue and then mounted with ProLong Diamond Antifade Mountant with DAPI (P36971, Thermo Fisher Scientific, Waltham, MA). Images were acquired using a Nikon A1 laser-scanning fluorescence confocal microscope. Photomultiplier tube (PMT) gain and offset were adjusted to obtain an optimal signal-to-noise ratio. Seven to ten images per sample were analyzed to measure the mean fluorescence intensity per unit area.

### RNAscope

Detection of specific RNAs was performed by RNAscope *in situ* hybridization (Wang et al. 2012) on FFPE skin tissue sections. All the pre-treatment solutions and detection kits were purchased from ACD (Advanced Cell Diagnostics, Inc., Hayward, CA, USA). The instructions on RNAscope™ Multiplex Fluorescent Reagent Kit v2 document from the company were followed for the entire RNAscope process with minor adjustments. A standard oven from Thermo Scientific (PR305145G) was used for slide baking (60°C) and all incubation steps (40°C). Different probes were used as necessary for the available channels. The RNA-specific double z-probes included: Cyp1a1 (464611-C2), Cyp1b1 (412881-C3), Prdm1 (441871, encodes BLIMP1), Lrig1 (310521-C2), Myc (413451-C3), Scd1 (461641), Pparg (418821-C3). The probes were conjugated to TSA Vivid Fluorophore 520 (323271), TSA Vivid Fluorophore 570 (323272) or TSA Vivid Fluorophore 650 (323273). The fluorophores were diluted at 1:1500 in TSA buffer (322810).

In brief, 5 µm-thick FFPE tissue sections on Superfrost Plus microscope slides were baked at 60°C for 1 hour. Slides were then deparaffinized in xylene, followed by dehydration in a series of ethanol concentrations (100%, 95%, 75% and 50%). Hydrophobic barriers were drawn around the tissue sections using a hydrophobic pen (310018, Vector Laboratories, Newark, CA, USA). Pretreatment conditions were optimized according to the tissue type. Slides were first incubated for 10 minutes at RT with hydrogen peroxide, followed by a 2 minute DI water wash. The slides were incubated for 7 minutes in 1X target retrieval reagent at 95-99°C, followed by a 2 minute wash in DI water, and then transferred to 100% ethanol for 2 minutes. Slides were dried at RT. For the final pretreatment, slides were placed in a humidifying chamber and sufficient drops of Protease Plus were added to cover the entire tissue section. The slides were then incubated at 40°C for 25 minutes. After incubation, the slides were washed twice in DI water for 2 minutes each. Target probes were diluted together at 1:50 in either C1 probe or probe diluent (300041). The negative control slide was covered with the probe diluent, while the other slides were covered with enough drops of the target probe(s) solution to cover the tissue. Slides were incubated at 40°C for 2 hours and then washed twice in 1X wash buffer for 2 minutes and stored in 5X SSC buffer overnight. The next day, slides were washed twice in 1X wash buffer before continuing with the assay. Signals were developed separately for each channel: incubation with HRP C1 (or the respective channel) for 15 minutes, incubation with the fluorophore for 30 minutes, and with the HRP blocker for 15 minutes. All the incubation steps for signal development were performed at 40°C. Between each step, the slides were washed twice in 1X wash buffer. DAPI was then added to the tissue sections for 30 seconds, and the sections were mounted using either ProLong Diamond Antifade Mountant with DAPI (P36971) or ProLong Gold Antifade Mountant with DAPI (P36935) from Thermo Fisher Scientific, followed by coverslipping. The slides were stored in the dark at 4°C overnight to dry before capturing images. Fluorescence images were acquired using a Nikon A1 laser-scanning fluorescence confocal microscope equipped with a 40X immersion oil NA 1.0 objective and using a pinhole diameter of 1.2 Airy units. For each fluorophore channel, laser intensity was selected for producing the brightest fluorescence intensity signal with minimal saturation. Photomultiplier tube (PMT) offset and gain were set to values where the fluorescence background was minimal for negative control sections. All the settings for a given channel were applied uniformly to all samples within an experiment. At least 3 images containing well-structured hair follicles were captured from different regions of each skin section. In the representative images presented, the lookup table (LUT) values were sometimes adjusted equally in all images, to improve visualization of signals. When presenting a magnified view, the original Nikon image was resized in Photoshop to 300 dpi and cropped to a specific pixel size in the specific figure.

### Quantitation of RNAscope signals

#### i) Mean fluorescence intensity (MFI) and % area measurements

Mean fluorescence intensity refers to the sum of all pixel values within the area of interest divided by the number of pixels, where the value is normalized to µm² area. At least 3 images were analyzed for each tissue sample. First, spatial calibration was applied to obtain the results in µm², and global calibration was applied to all images. Next, an ideal threshold was selected which best represented the original 16-bit images. This was done by comparing the original 16-bit images while adjusting the threshold for 8-bit images to detect the dots. The same threshold was applied to all the images included in an analysis. Regions of interest (ROIs) were drawn for different regions to be analyzed (infundibulum, interfollicular epidermis, sebaceous gland). Measurements were set to calculate the mean gray value and area fraction. Area fraction (% area) represents the percentage of area occupied by a particular probe in a region, whereas average dots/positive cell (described below) is the number transcripts in a cell that expresses at least one transcript. These measurements show gene expression in a regional and cellular context.

#### ii) Average dots/positive cell analysis

At least 3 images were analyzed for each tissue sample. The ROIs were drawn on the DAPI image and then merged with other channels. The cell counter plugin of ImageJ was used to count the positive cells within the ROIs. The “analyze particles” tool was used to count the total number of particles (dots) in the respective ROI.

#### iii) Average dots/positive cell count from intensity

When clusters of dots were apparent, analysis of each transcript was performed according to ACD protocol (SOP #45-006). A total of 50 single dots were selected in tissue regions where transcripts were clearly expressed as single dots. The intensity was background corrected and the average intensity of a single dot was calculated according to the protocol. The total integrated density of an ROI was background corrected and divided by the average intensity per single dot to obtain an estimate of the total number of dots in the respective ROI. Finally, the estimated total dot number in a ROI was divided by the total number of positive cells in the ROI to get average dots/positive cells. Object and area calculations were all performed in ImageJ.

### Statistical analysis

Statistical analysis was performed using GraphPad Prism software version 10.5.0. For single comparisons, Student’s t-test was used to compare the means of two groups. For comparisons of multiple groups, one-way or two-way ANOVA followed by Tukey’s post-hoc or Fisher’s LSD tests were performed based on the number of relevant comparisons considered. *p*-value > 0.05 were considered not significant, ns, * *p* < 0.05, ** *p* < 0.01, *** *p* < 0.001, **** *p* < 0.0001.

## Results

### *Ahr-*dependent effects of TCDD on the sebaceous gland

*In utero* and lactational exposure to TCDD results in the largest effect on SG atrophy at P21 in murine skin (Bhuju et al. 2021). To determine if this effect was *Ahr*-dependent, tissue from *Ahr +/+* and *Ahr -/-* pups were examined in the current study. TCDD-mediated SG atrophy was prominent in treated *Ahr +/+* skin, as previously observed **(Fig. 1A)**. This effect was absent in *Ahr*-null skin, indicating *Ahr*-dependency of TCDD-mediated seboatrophy. Oil Red O staining further confirmed this effect of diminishing gland size **(Fig. S2)**. Quantitative analysis showed that the gland sizes in TCDD-treated skin were about 32% of the area of the glands in the control skin; this effect was not significant in the *Ahr* -/- samples **(Fig. 1B)**. No obvious differences in the hair cycle were observed between vehicle- and TCDD-treated skin.

**Figure 1.**
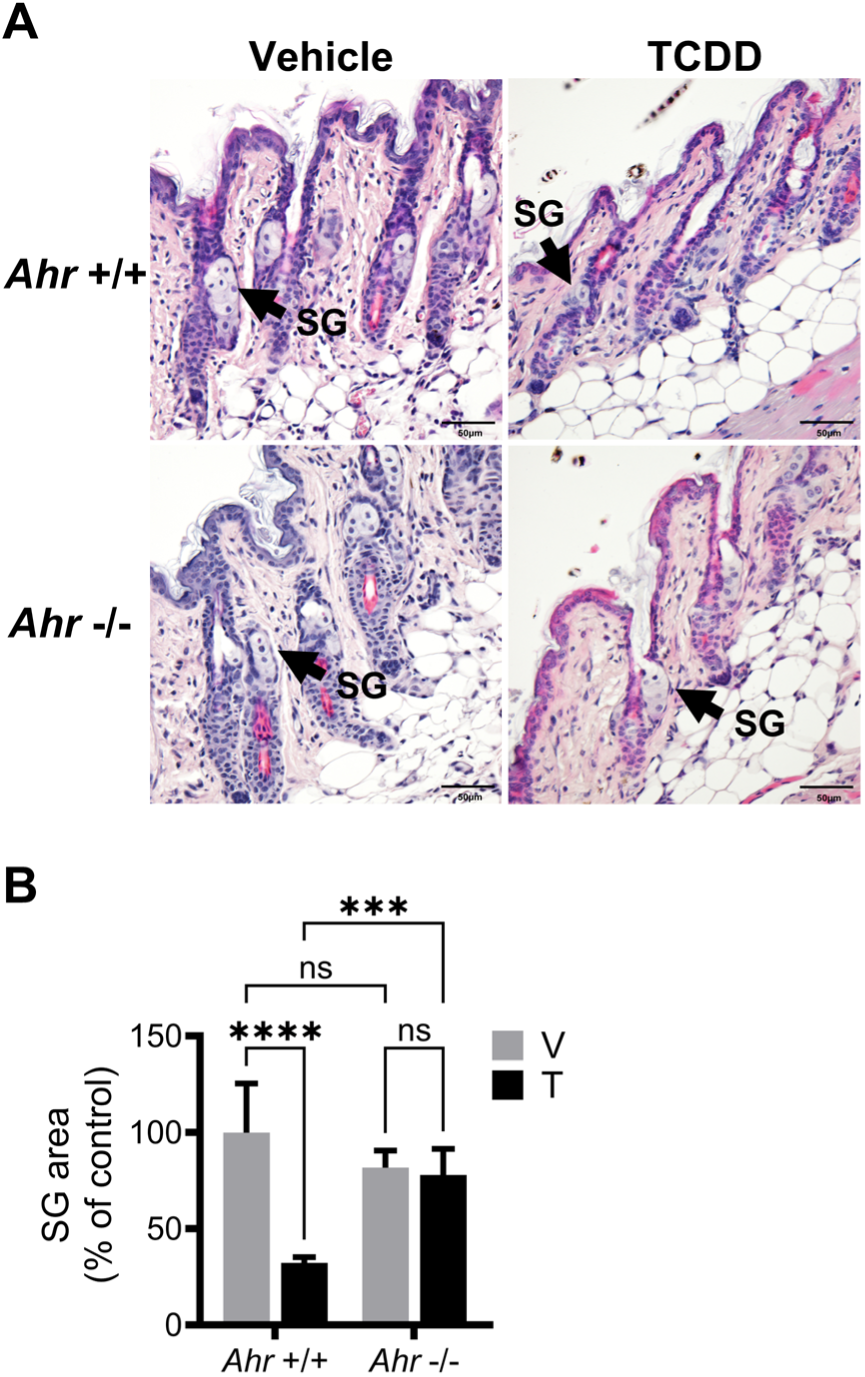
*Ahr*-dependence of ligand-induced sebaceous gland (SG) atrophy at P21. Time-mated heterozygous *Ahr* allele dams were treated with either vehicle (V, corn-oil) or TCDD (T, 5 µg/ kg bw) by oral gavage and *Ahr +/+* or *Ahr -/-* pup skin tissue (P21) was analyzed for SG effects. **(A)** Representative H&E skin images of vehicle- and TCDD-treated FFPE tissue sections at P21. Scale bar = 50 µm. Arrows point towards the SG. **(B)** SG area (mean ± SD, n = 5-6), is expressed as the percent of *Ahr +/+* vehicle (100%). Statistics were performed using a two-way ANOVA followed by Tukey’s multiple comparisons test, *** *p* < 0.001, **** *p* < 0.0001, ns, not significant.

### Cell- and region-specific responses to TCDD following *in utero* and lactational exposure

To determine the expression pattern and abundance of *Cyp1a1* RNA in the infundibulum (IF), RNAscope was performed on P21 mouse skin. Quantitative analyses included determination of the average number of RNA transcripts (dots) per positive cell and the area fraction occupied by positive cells (% area). The former measures cell specific levels of RNA; the latter estimates the percentage of transcript-expressing cells in a specific region of interest. Regional analysis of *Cyp1a1* in the IF showed that RNA expression was increased, correlating with previously reported effects on CYP1A1 protein, and that this increase was *Ahr*-dependent **(Fig. 2A-C and Fig. S3)**. The estimated number of transcripts per positive cell increased from around 2 per cell in vehicle-to 32 per cell in TCDD-treated *Ahr* +/+ skin. In the SG, some basal level of *Cyp1a1* was observed, independent of the *Ahr* genotype. The number of SG *Cyp1a1* transcripts per positive cell increased by about 9.5-fold in the treated +/+ group compared to the control and this effect was dependent on the *Ahr*. This metric of response is consistent with increased ligand-receptor occupancy in the responsive cells.

**Figure 2.**
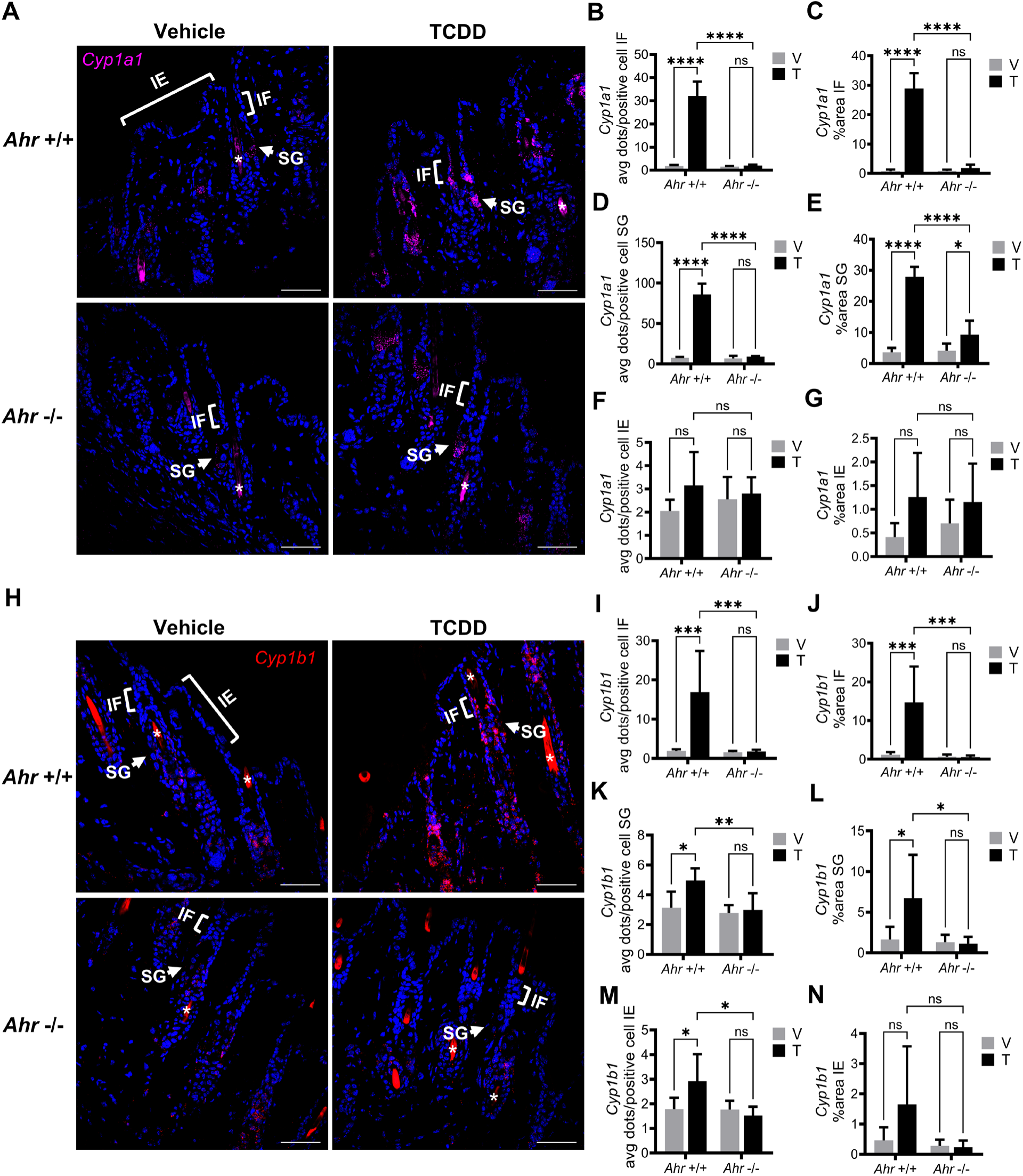
Single-cell and regional *in situ* analysis of *Cyp1a1* and *Cyp1b1* RNA at P21. Time-mated heterozygous *Ahr* allele dams were treated with either vehicle (V, corn-oil) or TCDD (T, 5 µg/ kg bw) by oral gavage and *Ahr +/+* or *Ahr -/-* pup skin tissue (P21) was analyzed. **(A)** Representative images of *Cyp1a1* RNA in murine skin. Lower infundibulum (IF), sebaceous gland (SG) and interfollicular epidermis (IE) are marked with brackets and arrows. Hair shaft is indicated by a white asterisk. Quantification of *Cyp1a1* in the IF by measuring **(B)** average dots per positive cell and **(C)** percent area. **(D-E)** Quantification of *Cyp1a1* in SGs; **(F-G)** in IE. **(H)** Representative images of *Cyp1b1* RNA in murine skin. Quantification of *Cyp1b1* in the IF by measuring **(I)** average dots per positive cell and **(J)** percent area. **(K-L)** Quantification of *Cyp1b1* in SGs; **(M-N)** in IE. Scale bar = 50 µm. Levels are expressed as mean ± SD (n = 5-6). Statistics were performed using a two-way ANOVA followed by Tukey’s multiple comparisons test, * *p* < 0.05, ** *p* < 0.01, *** *p* < 0.001, **** *p* < 0.0001, ns, not significant. No significant difference was identified for comparison of vehicle *Ahr* +/+ and vehicle *Ahr* -/- for any region.

The SG percent area of the total SG ROI also increased in response to treatment, with a notable increase of *Cyp1a1* in the lower cells of sebaceous glands **(Fig. 2D-E)**. Of note, a much smaller, yet significant increase of *Cyp1a1* percent area also was observed in the TCDD-treated *Ahr* -/-samples. No significant difference was found when comparing the control *Ahr* +/+ to control *Ahr -*/- percent areas, so this appears to be an effect of TCDD, independent of the *Ahr*. This effect could involve beta-catenin, as some evidence suggests that basal *Cyp1a1* expression is affected by a beta-catenin-dependent mechanism (Braeuning et al. 2011), and beta-catenin is known to play important roles in SG development and homeostasis (Geueke and Niemann 2021). Interestingly, the upper, more differentiated sebocytes barely expressed *Cyp1a1* under untreated conditions. *Cyp1a1* levels in the interfollicular epidermis (IE) were similar in vehicle- and TCDD-treated samples, independent of genotypes **(Fig. 2F-G and Fig. S4A)**.

The levels of *Cyp1b1* RNA, another biomarker of AHR response to ligand, were also measured. *Cyp1b1* expression was lower than *Cyp1a1*, but like *Cyp1a1* it was significantly elevated in the IF of TCDD-treated skin **(Fig. 2H-J)**. This effect was also *Ahr*-dependent. Estimated transcripts per positive cell increased from around 2 per cell in vehicle- to 17 per cell in TCDD-treated *Ahr* +/+ skin. While *Cyp1a1* occupied over 28% of the area of the IF, *Cyp1b1* occupied less than 15% of the same region. In SG, *Cyp1b1* levels were much lower compared to *Cyp1a1*, yet significantly elevated in terms of transcripts per cell and percent area **(Fig. 2K-L)**. Unlike *Cyp1a1*, *Cyp1b1* was increased in the IE by TCDD treatment in an *Ahr*-dependent manner **(Fig. 2M-N and Fig. S4B)**. In the IE, *Cyp1b1* transcripts ranged from less than 2 in the vehicle- to around 3 per positive cell in the TCDD-treated *Ahr* +/+ skin.

### Examining potential cell interactions of *Blimp1*, *Lrig1*, and *Myc* at P21

To explore a proposed mechanism linking TCDD-mediated SG atrophy and genes related to gland morphology, we analyzed the expression of *Blimp1*, *Lrig1*, and *Myc*. The prioritization of these transcripts was based on a model of sebocyte development that was originally proposed by Horsely et al. (Horsley et al. 2006). This model, later revised by Kretzschmar and colleagues (Kretzschmar et al. 2014), and then simplified by Bock (Bock 2016) is shown in **Figure 3A**. This model purports the existence of a bona fide SG stem cell population in the upper hair follicle and SG periphery and involves MYC in driving these stem cells from a quiescent state to a proliferative state associated with SG morphogenesis. BLIMP1 is proposed to repress *Myc* expression, thereby inhibiting proliferation (**Fig. 3A**). The seboatrophy observed in P21 TCDD-treated *Ahr* +/+ skin was associated with *Blimp1* induction in the lower IF/JZ **(Fig. 3B)** and this effect was *Ahr*-dependent **(Fig. 3B-D)**. In the IF and near the sebaceous duct cells (Kretzschmar et al. 2014), *Blimp1* was mainly expressed in the cells lining the inner surface of IF (Welle 2023), and occasionally colocalized with *Myc*. In this same region of the IF, *Lrig1* and *Myc* often colocalized and were primarily expressed in the basal cells of the outer surface adjacent to the basement membrane **(Fig. 3C).** This was true for skin from both vehicle- and TCDD-treated mice. In the sebaceous duct, *Blimp1* colocalized with *Lrig1* in specific cells (refer to **Fig. 7B, right,** for cell locations). The progression from infundibulum to SG formation included cells with expression of *Lrig1*+ *Blimp1*+, and *Lrig1*+ *Myc*+. This expression was not continuous in adjacent cells but rather interspersed along the basal epithelium of the gland. At the lower tip of the SG, transitional basal cells expressing high levels of *Myc* were observed, as previously reported (Cottle et al. 2013). Within the gland, transitional basal cells progressed to become sebocytes containing high *Myc* and lower *Blimp1*, and then differentiated further into the sebocytes in the outward part of the gland that express less *Myc* and higher *Blimp1*. Although *Blimp1* was induced in the lower infundibulum of treated *Ahr* +/+ skin, no changes were observed in *Myc* and *Lrig1* transcripts compared to control **(Fig. 3E-F)**, (for full images see **Fig. S5A-B)**. Therefore, the main difference between control and treated skin was the induction of *Blimp1*. However, *Blimp1* induction occurred infrequently in *Lrig1+* cells, suggesting that the mechanism of sebocyte differentiation is more complex than the current proposed model **(Fig. 3A)**.

**Figure 3.**
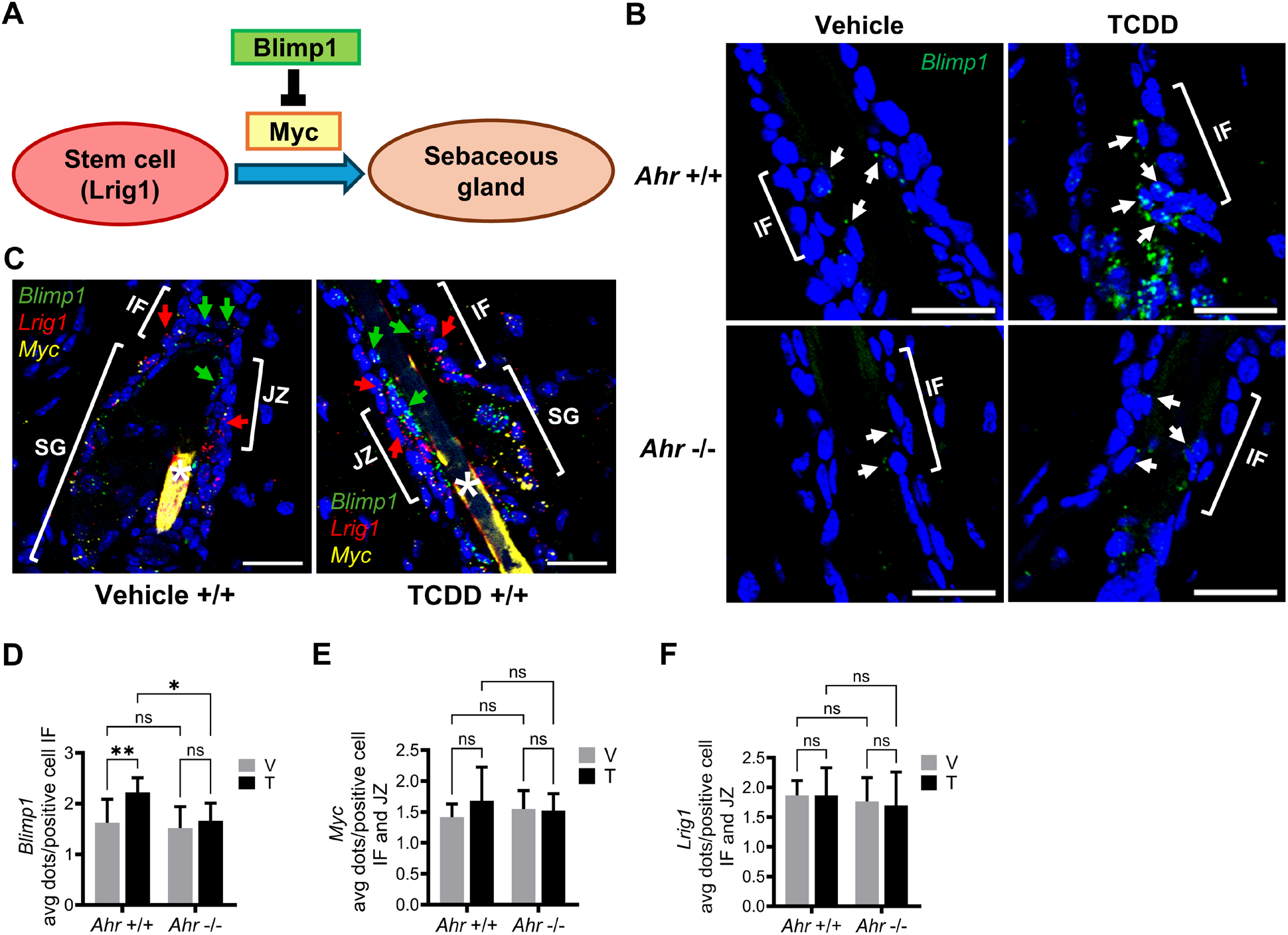
*Blimp1*, *Lrig1* and *Myc* expression in the pilosebaceous unit at P21. **(A)** A literature-based schema depicting a hypothetical role of *Blimp1* to affect differentiation of *Lrig1*+ progenitor cells during SG morphogenesis. **(B)** *Blimp1* expression in the lower infundibulum (IF) at P21. Time-mated mice were treated with either vehicle (corn oil) or TCDD (5 µg/ kg bw) via oral gavage and pups were harvested for analysis. Representative images (magnified view), with white arrows pointing towards cells expressing *Blimp1*. Scalebar = 20 µm. **(C)** *Blimp1*, *Myc* and *Lrig1* expression in vehicle- and TCDD-treated skin. Red arrows point towards the Lrig1-expressing cells in the basal cells of the outer surface and green arrows point towards the Blimp1-expressing cells lining the inner surface of the IF and JZ. Hair shaft background is identified by a white asterisk. **(D)** Quantification (average dots per positive cell) of *Blimp1* in the IF at P21. **(E)** Quantification of *Myc* average dots per positive cell in IF and JZ. **(F)** Quantification of *Lrig1* average dots per positive cell in IF and JZ at P21. The data is shown as mean ± SDs (n = 5-6). Statistics were performed using a two-way ANOVA. Subsequent analysis of the main effects was performed using the Fisher’s LSD test. ns, not significant, * *p* < 0.05, ** *p* < 0.01. LUT values were equally adjusted equally for all samples for ease of visualization.

### 3-Day TCDD dose-dependent induction of *Cyp1a1* and *Cyp1b1*

A 3-day dose-response study was conducted to evaluate biomarker response to TCDD following short-term topical exposure of the dorsal skin of adult mice. At the lowest dose (TCDD 0.0000003%), *Cyp1a1* expression was primarily increased in the cells of the lower infundibulum and the less differentiated sebocytes in the lower part of sebaceous gland. This expression increased in a dose-dependent manner **(Fig. 4A)**. Levels of transcripts in *Cyp1a1*-expressing cells were greater at each concentration of TCDD in the SG **(Fig. 4C)**. The IF and IE showed similar levels of transcripts **(Fig. 4B and Fig. 4D)**. The SG appeared slightly more sensitive to TCDD than the IE or IF, but the number of log doses used was too few to properly evaluate this parameter **(Fig. 4B-D)**. *Cyp1b1* showed regional expression and transcript levels that were similar to *Cyp1a1* **(Fig. 4E-H)**. Like *Cyp1a1*, *Cyp1b1* was more specifically expressed in the lower, less differentiated sebocytes, except at the highest concentration of TCDD, where this became less obvious. No sex-wise difference in any analysis of *Cyp1a1* or *Cyp1b1* was observed (data not shown).

**Figure 4.**
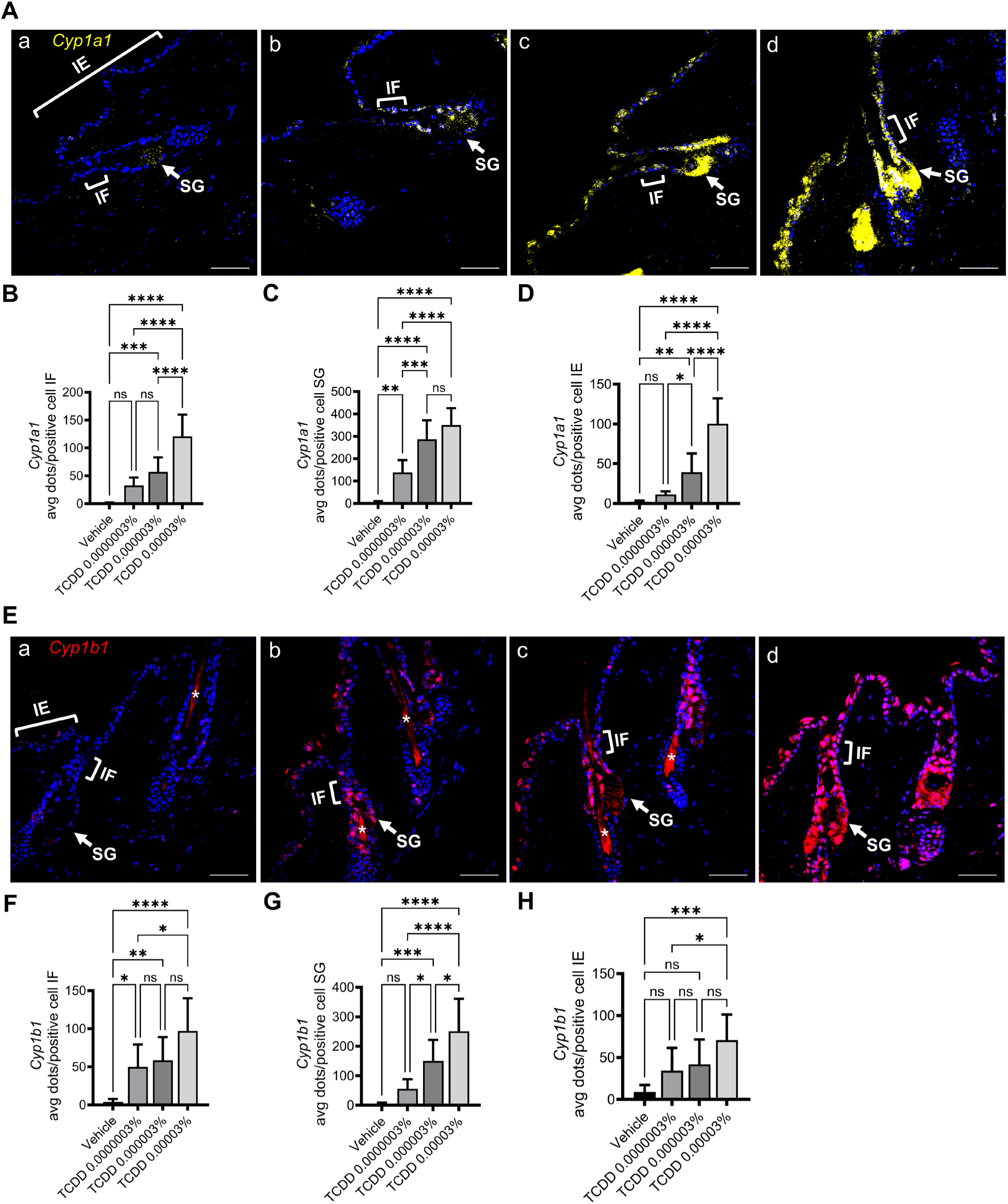
Dose-dependent induction of *Cyp1a1* and *Cyp1b1* following a 3-day topical exposure to vehicle [DMSO:acetone (1:9)] or TCDD at the indicated concentrations. **(A)** Representative images of *Cyp1a1* RNA expression in control and treated murine skin. (a) vehicle, (b) 0.0000003% TCDD, (c) 0.000003% TCDD, (d) 0.00003% TCDD. Lower infundibulum (IF), sebaceous gland (SG) and interfollicular epidermis (IE) are marked with brackets and arrows. Quantification of *Cyp1a1* expressed as average dots per positive cell in **(B)** IF; **(C)** SG; **(D)** IE. The data is shown as mean ± SDs (n = 8). Scale bar = 50 µm. **(E)** Representative images of *Cyp1b1* RNA expression in murine skin treated (a-d) as in (A). Quantification of *Cyp1b1* (average dots per positive cell) in **(F)** IF; **(G)** SG; **(H)** IE. Statistics were performed using a one-way ANOVA followed by Tukey’s multiple comparisons test, ns, not significant, * *p* < 0.05, ** *p* < 0.01, *** *p* < 0.001, **** *p* < 0.0001.

### Effects of 3-day TCDD treatment on sebocyte differentiation

Because of the dose-dependent and regional expression of *Cyp1a1* and *Cyp1b1* in the SG, we analyzed several endpoints of SG differentiation. The enzyme SCD1 (stearoyl-CoA desaturase) is known to have a role in sebocyte differentiation and is expressed in the sebaceous gland (Miyazaki et al. 2001; Zheng et al. 1999). Regional analysis of *Scd1* RNA expression in the SG was used to assess the effects of TCDD on early sebocyte differentiation. Co-expression of *Cyp1a1* with *Scd1* was used to determine the cellular and regional response to TCDD. Vehicle-treated skin exhibited very low expression of *Cyp1a1* **(Fig. 5Aa)**, whereas TCDD-treated (0.000003%) skin showed a robust increase in SG *Cyp1a1* transcript levels **(Fig. 5Ab)**. In the same samples, *Scd1* expression increased in percent area, with characteristics of a regional expansion that was more restricted than *Cyp1a1* **(Fig. 5Aa-b)**. Quantitative measurements of *Scd1* % area of the SG demonstrated a significant increase of regional *Scd1* area in the sebaceous glands of TCDD-treated mice, increasing from 40% in animals treated with vehicle to 50% in TCDD-treated samples **(Fig. 5B)**. For *Cyp1a1*, the area increased from less than 8% in vehicle- to over 62% in TCDD-treated samples **(Fig. 5C)**. As sebum production is linked to SG differentiation, Oil Red O staining was performed to assess production of lipids. Higher levels of Oil Red O staining were observed in the TCDD-treated samples **(Fig. 5D and Fig. S6)**. In addition, quantitation of SG lipid was performed using fluorescent Nile Red staining of the SG **(Fig. 5E and Fig. S7)**. Quantitative analysis of fluorescence intensity revealed a significant, 70%, increase in sebum production after 3 days of treatment with TCDD **(Fig. 5F)**. Neither Oil Red O **(Fig. S6C)** or Nile Red lipid staining **(Fig. S7C)** indicated a change in the SG area, supporting the idea of increased lipid production rather than increased gland size following a 3-day TCDD-treatment. PPARG is thought to play an important role in sebaceous gland formation and differentiation (Rosen et al. 1999; Veniaminova et al. 2023). However, no significant changes were observed in *Pparg* RNA expression following TCDD treatment **(Fig. S8)**.

**Figure 5.**
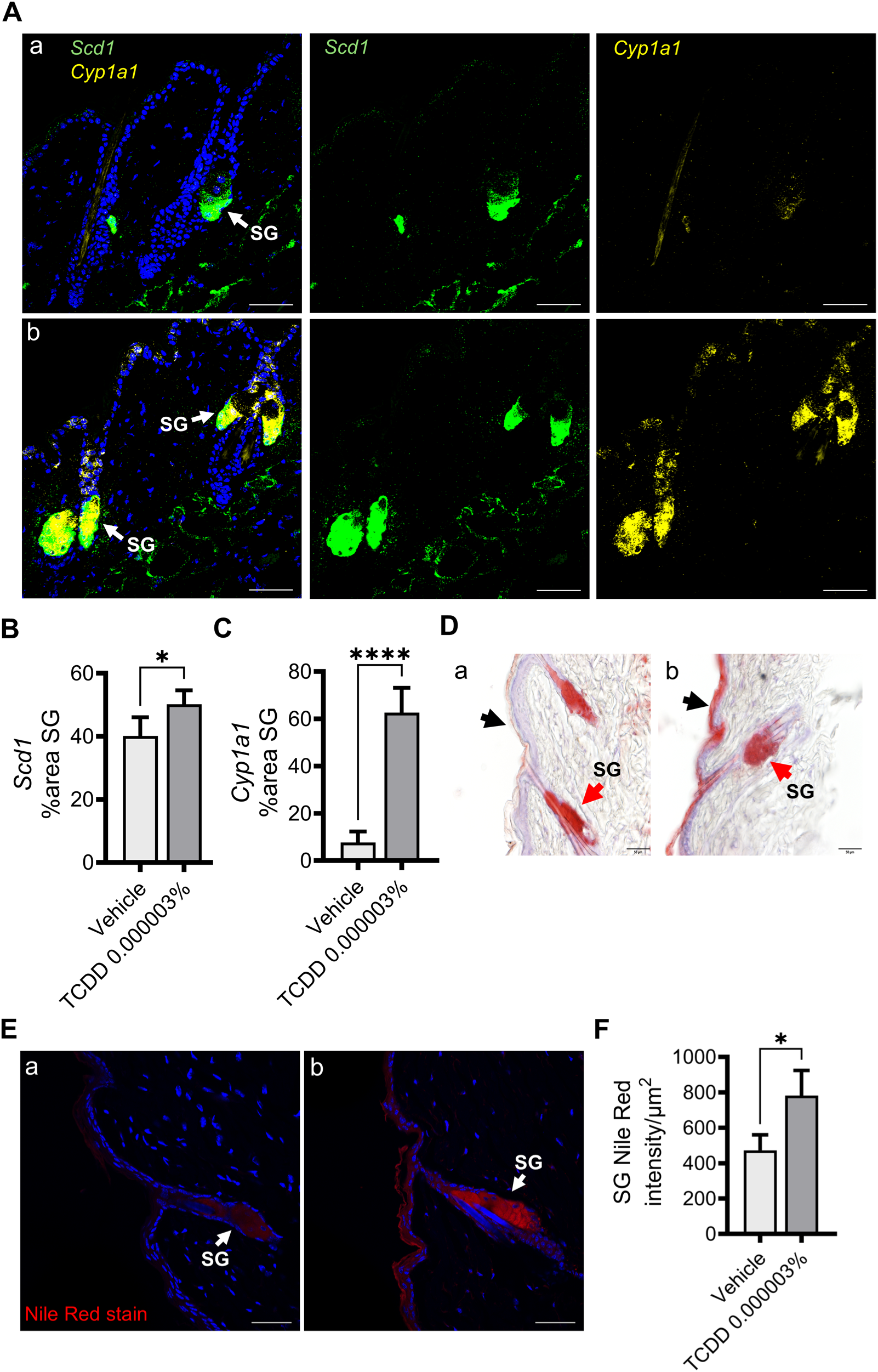
Early effect of TCDD on murine sebocyte differentiation and sebum production following 3-day topical exposure. **(A)** Representative images of *Scd1 and Cyp1a1* RNA expression in (a) vehicle or (b) 0.000003% TCDD treated skin. Arrows point towards SG. Scale bar = 50 µm. **(B)** Quantification of *Scd1* RNA in the SG, expressed as percent area. **(C)** Quantification of *Cyp1a1* RNA in the SG, expressed as percent area. The data (B-C) is shown as mean ± SDs (n = 5). Statistics were performed using a two-tailed Student’s t-test, * *p* < 0.05, **** p < 0.0001. **(D)** Representative images of Oil Red O staining in (a) vehicle- or (b) 0.000003% TCDD-treated skin. Arrows (black) point towards the interfollicular epidermis (IE); red arrows towards the SG. **(E)** Representative images of Nile Red lipid staining of (a) vehicle-or (b) 0.000003% TCDD-treated skin. **(F)** Quantitative analysis of fluorescence of Nile Red staining of SG lipids. The data is shown as mean ± SDs (n = 3) of intensity per unit area. Statistics were performed using a two-tailed Student’s t-test, * *p* < 0.05.

Observations of *Blimp1, Lrig1* and *Myc* RNA expression in 3-day vehicle- and TCDD-treated adult skin revealed a pattern of expression that is similar to the expression pattern of P21 mice in the lower infundibulum and sebaceous gland **(Fig. S5)**. A comparison of vehicle- and TCDD-treated skin showed that TCDD increased *Blimp1* in the IF and JZ, as described in P21 mice **(Fig. 6A and Fig. 6C)**. Previously, *Blimp1* was shown to be transcriptionally activated by AHR ligands in SZ95 cells (Ikuta et al. 2010). Like P21 mice, quantitative analysis confirmed the induction of *Blimp1* in the lower IF of TCDD-treated skin **(Fig. S9)**. *Blimp1* also appeared to be increased in the suprabasal layer of the epidermis **(Fig. 6A and Fig. 6C, blue arrow heads)**, marking keratinocyte terminal differentiation, as previously reported (Kretzschmar et al. 2014). In the SG of vehicle-treated skin, the lower sebocytes expressed higher *Myc* and little to no *Blimp1*, whereas more differentiated sebocytes expressed more *Blimp1* **(Fig. 6B)**. In TCDD-treated skin, we observed sebocytes in the outward gland to have higher numbers of *Blimp1* transcripts **(Fig. 6D)**. Quantitative analysis showed that the number of *Blimp1*+ sebocytes having more than 5 transcripts per cell increased from less than 2 in the vehicle-treated samples, to more than 3 in the sebocytes of the TCDD-treated skin **(Fig. 6E)**. This increase in *Blimp1+* staining sebocytes is consistent with increasing numbers of mature sebocytes (Kretzschmar et al. 2014). Overall, *Blimp1* appeared elevated in TCDD-treated skin, and accelerated differentiation of the sebaceous gland appears to be a major effect of TCDD on the pilosebaceous unit, along with increased differentiation of the epidermis.

**Figure 6.**
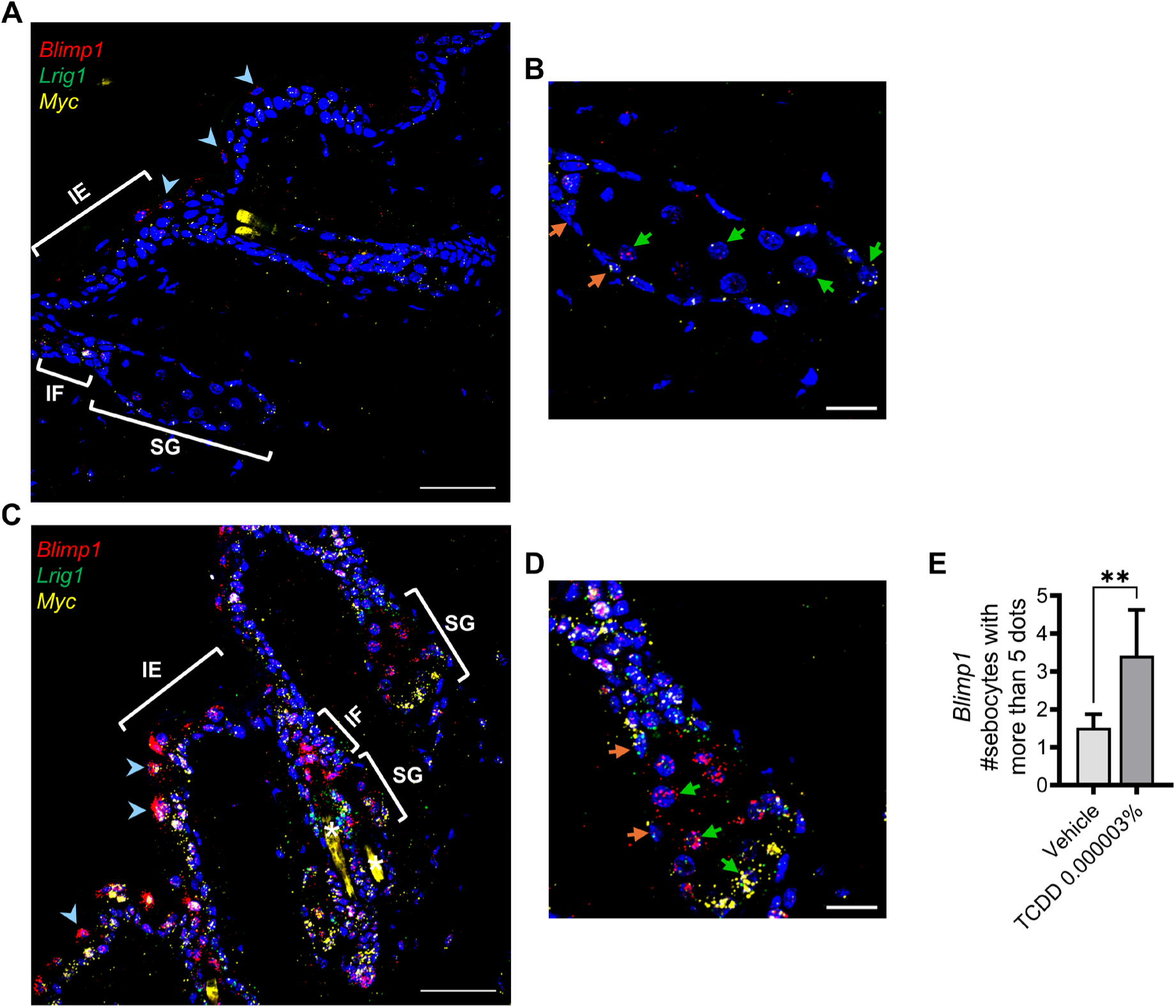
*Blimp1*, *Lrig1* and *Myc* RNA expression in the pilosebaceous unit in 3-day topical exposure. **(A and C)** Overview of *Blimp1, Lrig1, Myc* expression in skin treated topically for 3 days with either **(A-B)** vehicle (DMSO:acetone, 1:9) or **(C-D)** TCDD 0.000003%. Blue arrows point towards certain *Blimp1* expressing cells in the suprabasal layer of the IE. Scalebar = 50 µm. The hair shaft background is identified by a white asterisk. **(B and D)** Expression of *Blimp1*, *Lrig1*, *Myc* signals in SGs from vehicle- and TCDD-treated skin. Orange arrows point toward SG basal epithelial cells and green arrows point toward sebocytes with the following progression: high *Myc*, low *Blimp1* in the lower sebocytes to low *Myc*, high *Blimp1* in the upper sebocytes. Images are a magnified view. Scalebar = 20 µm. **(E)** Quantification of the number of sebocytes with more than 5 dots per cell of *Blimp1*. The data is shown as mean ± SDs (n = 5). Statistics were performed using a two-tailed Student’s t-test. ** *p* < 0.01. LUT values were equally adjusted for ease of visualization.

## Discussion

The sebaceous gland is an exocrine gland that produces sebum, a lipid containing substance that protects hair and enhances epidermal barrier function (Jang et al. 2018). This gland is a primary target of TCDD in humans, and SG atrophy and the formation of benign hamartomas along with hyperkeratosis of the skin are major effects in exposed people (Saurat et al. 2012). A similar, chloracne-like effect has been reported in mice exposed during development or topically to TCDD (Bhuju et al. 2021; Fontao et al. 2018). We previously reported that *in utero* and lactational exposure of mice to TCDD causes the highest SG atrophy at P21 of exposure (Bhuju et al. 2021). Our current findings show that this TCDD-mediated seboatrophy is dependent on the *Ahr*, like most other toxic effects of TCDD (Fernandez-Salguero et al. 1996).

One study of chloracne in mice reports the co-localization of TCDD-inducible CYP1A1 and LRIG1 in the lower infundibulum/junctional zone of the hair follicle where the LRIG1+ stem cells reside. The authors propose that TCDD targets these cells to induce SG atrophy (Fontao et al. 2018). Comparing *Cyp1a1* and *Cyp1b1* transcript levels in response to TCDD, we observed preferential TCDD-mediated induction of *Cyp1a1* in the IF/JZ and lower SG whereas *Cyp1b1* expression was induced throughout the epidermis, IF and SG. This confirms the results of Fontao and colleagues at the level of RNA and further demonstrates that different regions and cells of the epidermis and pilosebaceous unit selectively respond to TCDD-activated AHR.

According to current understanding, the LRIG1+ stem cells are bipotential during homeostasis (Jensen et al. 2009) and contribute to the maintenance of both the IF and SG (Geueke and Niemann 2021) The requirement of this LRIG1+ stem cell population for SG maintenance is firmly established in a recent study reporting that inducible-targeted ablation of LRIG1+ stem cells causes complete SG loss (Barnes et al. 2026). MYC regulates both cell fate and proliferation of the LRIG1+ cells to form the basal epithelium of the SG as well as the differentiation of the transitional basal cells into sebocytes (Cottle et al. 2013; Niemann and Horsley 2012). BLIMP1 is a repressor of MYC, and loss of *Blimp1* results in overexpression of MYC accompanied by SG hypertrophy and an increase of proliferative cells in the SG (Horsley et al. 2006). Thus, the interplay of BLIMP1 and MYC is believed to regulate SG homeostasis.

To explore the potential interactions of these known regulators of SG morphogenesis and maintenance we analyzed *Blimp1*, *Myc* and *Lrig1* transcripts in the skin of P21 pups, where TCDD-induced seboatrophy was observed. Levels of *Blimp1* transcripts were increased by TCDD in the lower infundibulum. This increase was dependent upon the *Ahr*, consistent with *in vitro* studies reporting that *Blimp1* expression is transcriptionally regulated by the ligand-activated AHR (Ikuta et al. 2010). However, the paradigm that BLIMP1 represses MYC to inhibit the transition of quiescent LRIG1+ stem cells to a proliferative state proved too simple at the cellular level. Firstly, *Lrig1* and *Myc* were primarily expressed in the outer basal cell layer, whereas *Blimp1* expression was mostly restricted to the epithelial cells lining the inner surface of the lower IF (see schema, **Fig. 7A**). Secondly, the expression of neither *Myc* nor *Lrig1* was altered by TCDD. Still, *Blimp1* was occasionally observed in certain *Lrig1-*expressing sebaceous duct cells, pointing to an interaction of BLIMP1 and LRIG1 in this restricted cell target. Alternatively, the cell localization of *Blimp1* to the epithelial cells lining the inner surface of the IF and *Lrig1* to the outer basal cells may indicate a cell-cell signaling mechanism such as paracrine growth factor-receptor interactions to regulate LRIG1+ stem cell fate. An example of BLIMP1 in this role occurs in breast cancer, where expression of *Blimp1* affects epithelial-to-mesenchymal transition and cell invasiveness through complex regulation of TGF-beta signaling (Romagnoli et al. 2012). Moreover, TGF-beta is reported to impact both hair follicle stem cell populations and lipid biosynthesis in the SG (Lin and Yang 2013; McNairn et al. 2013), and more studies are needed to evaluate the potential contributions of TGF-beta to SG homeostasis.

**Figure 7.**
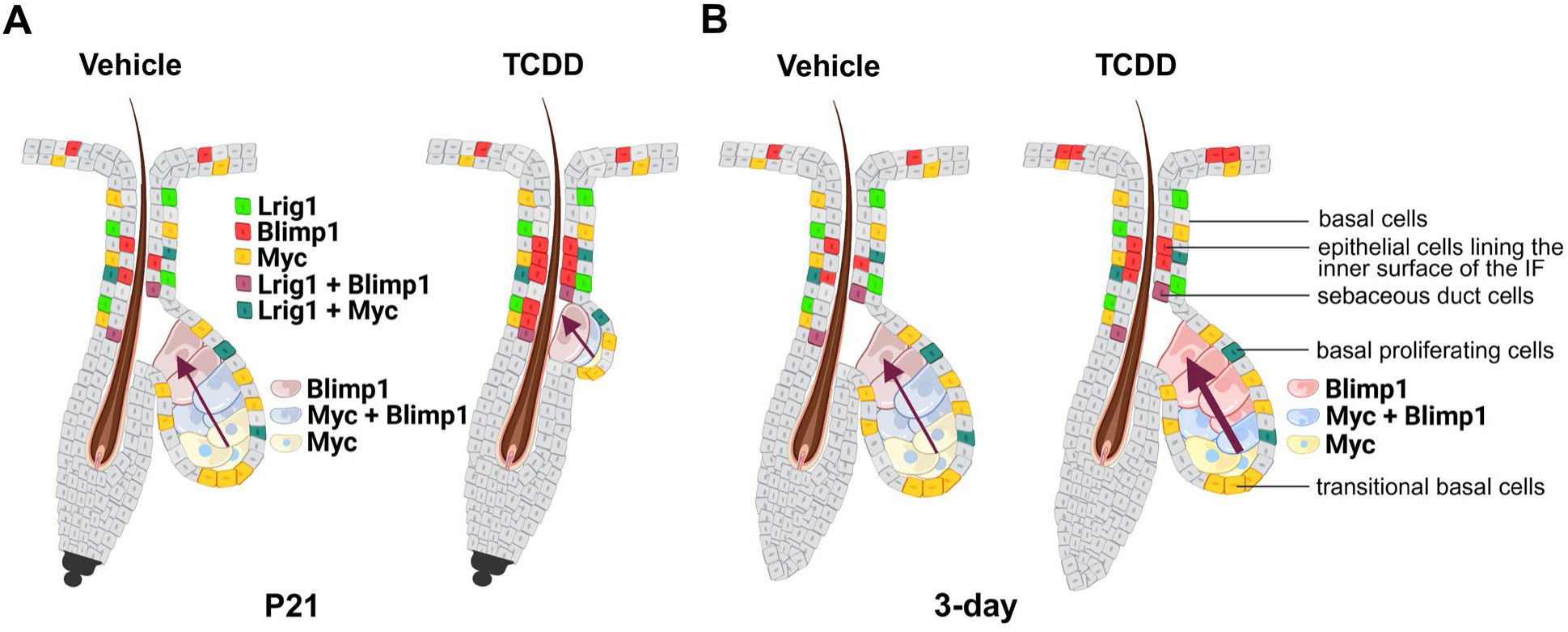
Summary schema of observed *Lrig1*, *Blimp1* and *Myc* expression in the lower infundibulum and sebaceous gland of *Ahr* +/+ murine skin in P21 and 3-day studies. **(A)** *Lrig1*, *Blimp1* and *Myc* expression in vehicle- and TCDD-treated skin at P21. **(Left)** In the lower infundibulum of control skin, the expression of *Lrig1* and *Myc* was primarily in the basal cells of the outer surface adjacent to the basement membrane and often colocalized; *Blimp1* expression was in the cells lining the inner surface; colocalized *Lrig1* and *Blimp1* expression was sometimes observed in specific sebaceous duct cells. In the sebaceous gland, *Myc* expression was in both the basal and the transitional basal cells, with higher expression in the transitional basal cells. In the sebocytes, *Myc* expression was highest in the lower, less differentiated cells, and decreased in the upper differentiated cells. *Blimp1* expression was found in the central sebocytes, with *Myc* and *Blimp1* expression often colocalized. **(Right)** In the lower infundibulum of P21 TCDD-exposed skin, *Lrig1* and *Myc* expression remained unchanged; *Blimp1* expression was increased in the inner cell layer of the infundibulum by TCDD. *Lrig1* and *Blimp1* sometimes colocalized in specific sebaceous duct cells. In the sebaceous gland, the pattern of gene expression was similar to what was observed in control skin, but the glands were smaller. **(B)** *Lrig1*, *Blimp1* and *Myc* expression in vehicle- and TCDD-treated skin from the 3-day study. **(Left)** In control skin, *Lrig1*, *Blimp1* and *Myc* expression was similar to the skin of P21 control mice. **(Right)** In TCDD-treated skin, expression of *Blimp1* is increased in the lower infundibulum and sometimes colocalized with *Lrig1* in the sebaceous duct cells. In the sebaceous gland, *Myc* levels appeared to be increased in the transitional basal cells. In sebocytes, TCDD samples showed fewer less centrally located sebocytes with coexpressed *Myc* and *Blimp1*. The number of sebocytes expressing higher levels of *Blimp1* transcripts was increased. Arrows show the direction of sebocyte differentiation. The thicker arrow represents increased differentiation. **Note:** Cell nomenclature was followed by literature: Basal cells (Veniaminova et al. 2023); Epithelial cells lining the inner surface of the IF (Welle 2023); Sebaceous duct cells (Kretzschmar et al. 2014); Basal proliferating cells (Kretzschmar et al. 2014); Transitional basal cells (Veniaminova et al. 2023).

In the SG, MYC plays a role in regulating sebocyte differentiation (Cottle et al. 2013). Sebocytes located at the lower tip of the SG express high levels of *Myc* and mark the transition from a basal proliferating cell to an early differentiating sebocyte. Within the gland, the sebocytes differentiate as they move inward and outward toward the duct canal, with less differentiating sebocytes expressing *Myc* and less *Blimp1*, and more highly differentiated sebocytes expressing only *Blimp1* (see schema, **Fig. 7A**) (Cottle et al. 2013; Kretzschmar et al. 2014). Although the TCDD-treated *Ahr* +/+ skin exhibited SG atrophy and less sebocytes, the above expression pattern of *Myc* and *Blimp1* transcripts was similar in control and treated samples, suggesting that TCDD alters the rate of sebocyte loss but not the mechanism of differentiation. Overall, the observed pattern of *Blimp1* expression suggests that BLIMP1 may contribute to cell state transitions during differentiation, rather than direct regulation of LRIG+ stem cell proliferation. Moreover, this concept is consistent with the role of BLIMP1 in guiding cell fate of effector B and T cells (Nutt et al. 2007) and plasma cell differentiation (Shaffer et al. 2002), as both cell fate decisions have been reported to be altered by TCDD (Schneider et al. 2009) and the *Ahr* (Vaidyanathan et al. 2017).

Because the preferential TCDD dose-dependent expression of *Cyp1a1* in the lower SG followed a pattern similar to SG differentiation, we analyzed several endpoints of this process *in vivo*. *Cyp1a1* colocalized with *Scd1* in the lower SG, and the area of *Scd1* expression significantly increased with TCDD treatment. SCD1 catalyzes the conversion of saturated fatty acids to monounsaturated fatty acids to affect SG lipid metabolism (Miyazaki et al. 2001; Zheng et al. 1999). Pharmacological inhibition of SCD1 (Meingassner et al. 2013) results in SG atrophy, and mutation (Sundberg et al. 2000) or targeted deletion of *Scd1* in the LRIG1 progenitor cell compartment (Lim et al. 2023) results in loss of hair and SG in the hair follicle. In the liver, *Scd1* is regulated by the *Ahr*, and *Scd1* null mice are resistant to TCDD-induced changes in lipid composition, hepatotoxicity, and triglyceride accumulation during the induction of steatosis (Angrish et al. 2011). In addition to expanding the *Scd1* area, Nile Red lipid staining showed a TCDD-dependent quantitative increase in SG lipids, occurring in parallel to the increased number of sebocytes progressing towards terminal differentiation. Enhanced sebocyte differentiation was further confirmed by analysis of the number of high *Blimp1*-expressing sebocytes that were located towards the outer SG duct canal (see schema, **Fig. 7B**).

Elevated expression of BLIMP1 in the suprabasal layers of the epidermis indicates keratinocyte differentiation (Chang et al. 2002; Lo Celso et al. 2008; Magnúsdóttir et al. 2007). Here we demonstrate that TCDD-mediated sebocyte differentiation parallels an increase in keratinocyte differentiation as evidenced by elevated *Blimp1* expression in the suprabasal epidermis. This is consistent with our previous reports of the effects of TCDD to accelerate epidermal barrier development and keratinocyte differentiation (Kennedy et al. 2013; Sutter et al. 2011). Of interest, metabolic reprogramming appears to regulate the differentiation of both sebocytes and keratinocytes. A recent study reports that sebaceous gland differentiation is governed, in part, by reprogramming of metabolic pathways involved in glycolysis, the TCA cycle, and redox metabolism (Schmidt et al. 2025). Remarkable similarities have been reported for the effects of TCDD to promote AHR-dependent keratinocyte differentiation by metabolic reprogramming that alters glycolysis and increases the cellular oxidative environment (Sutter et al. 2019).

Previously, an *in vitro* study demonstrated that TCDD alters differentiation in SZ95 sebocytes (Ju et al. 2011). Treatment of SZ95 cells with TCDD showed reduced levels of intracellular lipid. While these results seem contradictory to our *in vivo* results, it may reflect the plasticity of SZ95 cells to become either lipid producing sebocytes, or to differentiate into a K10 expressing keratinocytes. Such a bipotential generation of progeny having either phenotype has been described for the LRIG1+ progenitor cells (Jensen et al. 2009). Therefore, the differences in the effects of TCDD, *in vitro* versus *in vivo*, may be due to additional regulatory signals present *in vivo*, for example, cell-matrix, cell-adhesion, and growth factor signaling mechanisms. In P21 samples, protein expression of K10 was observed in the sebaceous duct cells in 4/5 TCDD-versus 1/5 vehicle-treated samples (data not shown). This staining pattern was virtually absent in 3-day TCDD-treated skin, suggesting that the SG ductal keratinization occurs later in response to exposure. While rarely occurring, this low frequency is consistent with our previous report of keratinous cyst-like structures that also occurred at low frequency in P21 TCDD-treated mice (Bhuju et al. 2021). More study will be required to determine whether our observations of ductal K10 expression, infrequent formation of keratinous cysts, and the trans-differentiation of SZ95 sebocytes to differentiated keratinocytes previously reported (Ju et al. 2011) are cellularly or mechanistically related.

Overall, the observed patterns of *Lrig1, Myc*, and *Blimp1* expression appear similar in the P21 and 3-day TCDD exposures, with only *Blimp1* showing a significant increase in response to TCDD in both the 3-day and P21 exposure studies. *In vivo*, the major effect of TCDD appears to be the early acceleration of SG differentiation with lipid production, followed by *Ahr*-dependent seboatrophy. Whether the regulation of SG maintenance is altered in response to TCDD remains equivocal, and additional studies using genetic modification and lineage tracing will be required. The effects of TCDD to accelerate SG and epidermal differentiation, accompanied by loss of sebocytes and enhanced epithelial keratinization, are strongly supported. Moreover, these effects may be sufficient to cause the resulting chloracne-like phenotypes and thus provide new insight into the etiology of chloracne.

## Data availability

All data are presented in the manuscript and the supplementary document. Additional information may be obtained upon request.

## Supplementary data

Supplementary material is available at *Toxicological Sciences* online.

## Acknowledgements

We would like to thank the Integrated Microscopy Center, University of Memphis, for the use of microscopes and histology. We are also grateful to Dr. Amy Abell, University of Memphis, for assistance with figure preparation. Finally, we want to thank Dr. Madhusudhan Alle for standardizing the Oil Red O stain protocol.

## References

Angrish MM, Jones AD, Harkema JR, Zacharewski TR. 2011. Aryl hydrocarbon receptor-mediated induction of stearoyl-coa desaturase 1 alters hepatic fatty acid composition in tcdd-elicited steatosis. Toxicol Sci. 124(2):299–310. doi: 10.1093/toxsci/kfr226.

Arnold I, Watt FM. 2001. C-myc activation in transgenic mouse epidermis results in mobilization of stem cells and differentiation of their progeny. Curr Biol. 11(8):558–568. doi: 10.1016/s0960-9822(01)00154-3.

Barnes L, Fontao F, Konstantinou E, Saurat JH, Sorg O, Kaya G. 2026. Specific in vivo ablation of lrig1-positive follicular progenitor cells results in sebaceous gland loss in mice. Int J Mol Sci. 27(3). doi: 10.3390/ijms27031513.

Bhuju J, Olesen KM, Muenyi CS, Patel TS, Read RW, Thompson L, Skalli O, Zheng Q, Grice EA, Sutter CH, Sutter TR. 2021. Cutaneous effects of in utero and lactational exposure of c57bl/6j mice to 2,3,7,8-tetrachlorodibenzo-p-dioxin. Toxics. 9(8). doi: 10.3390/toxics9080192.

Birnbaum LS. 1994. The mechanism of dioxin toxicity: Relationship to risk assessment. Environ Health Perspect. 102 Suppl 9(Suppl 9):157–167. doi: 10.1289/ehp.94102s9157.

Bock KW. 2016. Toward elucidation of dioxin-mediated chloracne and ah receptor functions. Biochem Pharmacol. 112(1–5. doi: 10.1016/j.bcp.2016.01.010.

Braeuning A, Köhle C, Buchmann A, Schwarz M. 2011. Coordinate regulation of cytochrome p450 1a1 expression in mouse liver by the aryl hydrocarbon receptor and the beta-catenin pathway. Toxicol Sci. 122(1):16–25. doi: 10.1093/toxsci/kfr080.

Chang DH, Cattoretti G, Calame KL. 2002. The dynamic expression pattern of b lymphocyte induced maturation protein-1 (blimp-1) during mouse embryonic development. Mech Dev. 117(1-2):305–309. doi: 10.1016/s0925-4773(02)00189-2.

Cottle DL, Kretzschmar K, Schweiger PJ, Quist SR, Gollnick HP, Natsuga K, Aoyagi S, Watt FM. 2013. C-myc-induced sebaceous gland differentiation is controlled by an androgen receptor/p53 axis. Cell Rep. 3(2):427–441. doi: 10.1016/j.celrep.2013.01.013.

Fernandez-Salguero PM, Hilbert DM, Rudikoff S, Ward JM, Gonzalez FJ. 1996. Aryl-hydrocarbon receptor-deficient mice are resistant to 2,3,7,8-tetrachlorodibenzo-p-dioxin-induced toxicity. Toxicol Appl Pharmacol. 140(1):173–179. doi: 10.1006/taap.1996.0210.

Fontao F, Barnes L, Kaya G, Saurat JH, Sorg O. 2018. High susceptibility of lrig1 sebaceous stem cells to tcdd in mice. Toxicol Sci. 161(1):207. doi: 10.1093/toxsci/kfx251.

Frances D, Niemann C. 2012. Stem cell dynamics in sebaceous gland morphogenesis in mouse skin. Dev Biol. 363(1):138–146. doi: 10.1016/j.ydbio.2011.12.028.

Geueke A, Niemann C. 2021. Stem and progenitor cells in sebaceous gland development, homeostasis and pathologies. Exp Dermatol. 30(4):588–597. doi: 10.1111/exd.14303.

Honeycutt KA, Roop DR. 2004. C-myc and epidermal stem cell fate determination. J Dermatol. 31(5):368–375. doi: 10.1111/j.1346-8138.2004.tb00687.x.

Horsley V, O’Carroll D, Tooze R, Ohinata Y, Saitou M, Obukhanych T, Nussenzweig M, Tarakhovsky A, Fuchs E. 2006. Blimp1 defines a progenitor population that governs cellular input to the sebaceous gland. Cell. 126(3):597–609. doi: 10.1016/j.cell.2006.06.048.

Hwa C, Bauer EA, Cohen DE. 2011. Skin biology. Dermatol Ther. 24(5):464–470. doi: 10.1111/j.1529-8019.2012.01460.x.

Ikuta T, Ohba M, Zouboulis CC, Fujii-Kuriyama Y, Kawajiri K. 2010. B lymphocyte-induced maturation protein 1 is a novel target gene of aryl hydrocarbon receptor. J Dermatol Sci. 58(3):211–216. doi: 10.1016/j.jdermsci.2010.04.003.

Jang H, Myung H, Lee J, Myung JK, Jang WS, Lee SJ, Bae CH, Kim H, Park S, Shim S. 2018. Impaired skin barrier due to sebaceous gland atrophy in the latent stage of radiation-induced skin injury: Application of non-invasive diagnostic methods. Int J Mol Sci. 19(1). doi: 10.3390/ijms19010185.

Jensen KB, Collins CA, Nascimento E, Tan DW, Frye M, Itami S, Watt FM. 2009. Lrig1 expression defines a distinct multipotent stem cell population in mammalian epidermis. Cell Stem Cell. 4(5):427–439. doi: 10.1016/j.stem.2009.04.014.

Ju Q, Fimmel S, Hinz N, Stahlmann R, Xia L, Zouboulis CC. 2011. 2,3,7,8-tetrachlorodibenzo-p-dioxin alters sebaceous gland cell differentiation in vitro. Exp Dermatol. 20(4):320–325. doi: 10.1111/j.1600-0625.2010.01204.x.

Kennedy LH, Sutter CH, Leon Carrion S, Tran QT, Bodreddigari S, Kensicki E, Mohney RP, Sutter TR. 2013. 2,3,7,8-tetrachlorodibenzo-p-dioxin-mediated production of reactive oxygen species is an essential step in the mechanism of action to accelerate human keratinocyte differentiation. Toxicol Sci. 132(1):235–249. doi: 10.1093/toxsci/kfs325.

Knutson JC, Poland A. 1982. Response of murine epidermis to 2,3,7,8-tetrachlorodibenzo-p-dioxin: Interaction of the ah and hr loci. Cell. 30(1):225–234. doi: 10.1016/0092-8674(82)90028-9.

Kretzschmar K, Cottle DL, Donati G, Chiang MF, Quist SR, Gollnick HP, Natsuga K, Lin KI, Watt FM. 2014. Blimp1 is required for postnatal epidermal homeostasis but does not define a sebaceous gland progenitor under steady-state conditions. Stem Cell Reports. 3(4):620–633. doi: 10.1016/j.stemcr.2014.08.007.

Lim SBH, Wei S, Tan AH, van Steensel MAM, Lim X. 2023. Lrig1-expressing epidermal progenitors require scd1 to maintain the dermal papilla niche. Sci Rep. 13(1):4027. doi: 10.1038/s41598-023-30411-7.

Lin HY, Yang LT. 2013. Differential response of epithelial stem cell populations in hair follicles to tgf-β signaling. Dev Biol. 373(2):394–406. doi: 10.1016/j.ydbio.2012.10.021.

Lo Celso C, Berta MA, Braun KM, Frye M, Lyle S, Zouboulis CC, Watt FM. 2008. Characterization of bipotential epidermal progenitors derived from human sebaceous gland: Contrasting roles of c-myc and beta-catenin. Stem Cells. 26(5):1241–1252. doi: 10.1634/stemcells.2007-0651.

Magnúsdóttir E, Kalachikov S, Mizukoshi K, Savitsky D, Ishida-Yamamoto A, Panteleyev AA, Calame K. 2007. Epidermal terminal differentiation depends on b lymphocyte-induced maturation protein-1. Proc Natl Acad Sci U S A. 104(38):14988–14993. doi: 10.1073/pnas.0707323104.

McNairn AJ, Doucet Y, Demaude J, Brusadelli M, Gordon CB, Uribe-Rivera A, Lambert PF, Bouez C, Breton L, Guasch G. 2013. Tgfβ signaling regulates lipogenesis in human sebaceous glands cells. BMC Dermatol. 13(2. doi: 10.1186/1471-5945-13-2.

Meingassner JG, Aschauer H, Winiski AP, Dales N, Yowe D, Winther MD, Zhang Z, Stütz A, Billich A. 2013. Pharmacological inhibition of stearoyl coa desaturase in the skin induces atrophy of the sebaceous glands. J Invest Dermatol. 133(8):2091–2094. doi: 10.1038/jid.2013.89.

Miyazaki M, Man WC, Ntambi JM. 2001. Targeted disruption of stearoyl-coa desaturase1 gene in mice causes atrophy of sebaceous and meibomian glands and depletion of wax esters in the eyelid. J Nutr. 131(9):2260–2268. doi: 10.1093/jn/131.9.2260.

Muenyi CS, Carrion SL, Jones LA, Kennedy LH, Slominski AT, Sutter CH, Sutter TR. 2014. Effects of in utero exposure of c57bl/6j mice to 2,3,7,8-tetrachlorodibenzo-p-dioxin on epidermal permeability barrier development and function. Environ Health Perspect. 122(10):1052–1058. doi: 10.1289/ehp.1308045.

Muku GE, Blazanin N, Dong F, Smith PB, Thiboutot D, Gowda K, Amin S, Murray IA, Perdew GH. 2019. Selective ah receptor ligands mediate enhanced srebp1 proteolysis to restrict lipogenesis in sebocytes. Toxicol Sci. 171(1):146–158. doi: 10.1093/toxsci/kfz140.

Niemann C. 2009. Differentiation of the sebaceous gland. Dermato-Endocrinology. 1(2):64–67. https://doi.org/10.4161/derm.1.2.8486. doi: 10.4161/derm.1.2.8486.

Niemann C, Horsley V. 2012. Development and homeostasis of the sebaceous gland. Semin Cell Dev Biol. 23(8):928–936. doi: 10.1016/j.semcdb.2012.08.010.

Nutt SL, Fairfax KA, Kallies A. 2007. Blimp1 guides the fate of effector b and t cells. Nat Rev Immunol. 7(12):923–927. doi: 10.1038/nri2204.

Panteleyev AA, Bickers DR. 2006. Dioxin-induced chloracne--reconstructing the cellular and molecular mechanisms of a classic environmental disease. Exp Dermatol. 15(9):705–730. doi: 10.1111/j.1600-0625.2006.00476.x.

Puhvel SM, Sakamoto M, Ertl DC, Reisner RM. 1982. Hairless mice as models for chloracne: A study of cutaneous changes induced by topical application of established chloracnegens. Toxicol Appl Pharmacol. 64(3):492–503. doi: 10.1016/0041-008x(82)90247-2.

Romagnoli M, Belguise K, Yu Z, Wang X, Landesman-Bollag E, Seldin DC, Chalbos D, Barillé-Nion S, Jézéquel P, Seldin ML, Sonenshein GE. 2012. Epithelial-to-mesenchymal transition induced by tgf-β1 is mediated by blimp-1-dependent repression of bmp-5. Cancer Res. 72(23):6268–6278. doi: 10.1158/0008-5472.Can-12-2270.

Rosen ED, Sarraf P, Troy AE, Bradwin G, Moore K, Milstone DS, Spiegelman BM, Mortensen RM. 1999. Ppar gamma is required for the differentiation of adipose tissue in vivo and in vitro. Mol Cell. 4(4):611–617. doi: 10.1016/s1097-2765(00)80211-7.

Saurat JH, Kaya G, Saxer-Sekulic N, Pardo B, Becker M, Fontao L, Mottu F, Carraux P, Pham XC, Barde C, Fontao F, Zennegg M, Schmid P, Schaad O, Descombes P, Sorg O. 2012. The cutaneous lesions of dioxin exposure: Lessons from the poisoning of victor yushchenko. Toxicol Sci. 125(1):310–317. doi: 10.1093/toxsci/kfr223.

Schmidt JV, Su GH, Reddy JK, Simon MC, Bradfield CA. 1996. Characterization of a murine ahr null allele: Involvement of the ah receptor in hepatic growth and development. Proc Natl Acad Sci U S A. 93(13):6731–6736. doi: 10.1073/pnas.93.13.6731.

Schmidt M, Binder H, Schneider MR. 2025. The metabolic underpinnings of sebaceous lipogenesis. Commun Biol. 8(1):670. doi: 10.1038/s42003-025-08105-9.

Schneider D, Manzan MA, Yoo BS, Crawford RB, Kaminski N. 2009. Involvement of blimp-1 and ap-1 dysregulation in the 2,3,7,8-tetrachlorodibenzo-p-dioxin-mediated suppression of the igm response by b cells. Toxicol Sci. 108(2):377–388. doi: 10.1093/toxsci/kfp028.

Schneider MR, Paus R. 2010. Sebocytes, multifaceted epithelial cells: Lipid production and holocrine secretion. Int J Biochem Cell Biol. 42(2):181–185. doi: 10.1016/j.biocel.2009.11.017.

Shaffer AL, Lin KI, Kuo TC, Yu X, Hurt EM, Rosenwald A, Giltnane JM, Yang L, Zhao H, Calame K, Staudt LM. 2002. Blimp-1 orchestrates plasma cell differentiation by extinguishing the mature b cell gene expression program. Immunity. 17(1):51–62. doi: 10.1016/s1074-7613(02)00335-7.

Solanki S, Tasnim SM, Thompson LL, Alle M, Ward B, Slominski AT, Janjetovic Z, Jozwicka T, Erdmańska P, Minzaghi D, Sutter CH, Grice EA, Sutter TR. 2026. Comparative 28-day mouse study of topically applied aryl hydrocarbon receptor ligands: Microbiota-derived indoles, therapeutic tapinarof, pollutants 2,3,7,8-tetrachlorodibenzo-p-dioxin and diesel exhaust particles. Chem Biol Interact. 433(112066. doi: 10.1016/j.cbi.2026.112066.

Sundberg JP, Boggess D, Sundberg BA, Eilertsen K, Parimoo S, Filippi M, Stenn K. 2000. Asebia-2j (scd1(ab2j)): A new allele and a model for scarring alopecia. Am J Pathol. 156(6):2067–2075. doi: 10.1016/s0002-9440(10)65078-x.

Suskind RR. 1985. Chloracne, “the hallmark of dioxin intoxication”. Scand J Work Environ Health. 11(3 Spec No):165–171. doi: 10.5271/sjweh.2240.

Sutter CH, Bodreddigari S, Campion C, Wible RS, Sutter TR. 2011. 2,3,7,8-tetrachlorodibenzo-p-dioxin increases the expression of genes in the human epidermal differentiation complex and accelerates epidermal barrier formation. Toxicol Sci. 124(1):128–137. doi: 10.1093/toxsci/kfr205.

Sutter CH, Olesen KM, Bhuju J, Guo Z, Sutter TR. 2019. Ahr regulates metabolic reprogramming to promote sirt1-dependent keratinocyte differentiation. J Invest Dermatol. 139(4):818–826. doi: 10.1016/j.jid.2018.10.019.

Vaidyanathan B, Chaudhry A, Yewdell WT, Angeletti D, Yen WF, Wheatley AK, Bradfield CA, McDermott AB, Yewdell JW, Rudensky AY, Chaudhuri J. 2017. The aryl hydrocarbon receptor controls cell-fate decisions in b cells. J Exp Med. 214(1):197–208. doi: 10.1084/jem.20160789.

Veniaminova NA, Jia YY, Hartigan AM, Huyge TJ, Tsai SY, Grachtchouk M, Nakagawa S, Dlugosz AA, Atwood SX, Wong SY. 2023. Distinct mechanisms for sebaceous gland self-renewal and regeneration provide durability in response to injury. Cell Rep. 42(9):113121. doi: 10.1016/j.celrep.2023.113121.

Wang F, Flanagan J, Su N, Wang LC, Bui S, Nielson A, Wu X, Vo HT, Ma XJ, Luo Y. 2012. Rnascope: A novel in situ rna analysis platform for formalin-fixed, paraffin-embedded tissues. J Mol Diagn. 14(1):22–29. doi: 10.1016/j.jmoldx.2011.08.002.

Welle MM. 2023. Basic principles of hair follicle structure, morphogenesis, and regeneration. Vet Pathol. 60(6):732–747. doi: 10.1177/03009858231176561.

Wysocki AB. 1999. Skin anatomy, physiology, and pathophysiology. Nurs Clin North Am. 34(4):777–797, v. doi.

Zanet J, Pibre S, Jacquet C, Ramirez A, de Alborán IM, Gandarillas A. 2005. Endogenous myc controls mammalian epidermal cell size, hyperproliferation, endoreplication and stem cell amplification. J Cell Sci. 118(Pt 8):1693–1704. doi: 10.1242/jcs.02298.

Zheng Y, Eilertsen KJ, Ge L, Zhang L, Sundberg JP, Prouty SM, Stenn KS, Parimoo S. 1999. Scd1 is expressed in sebaceous glands and is disrupted in the asebia mouse. Nat Genet. 23(3):268–270. doi: 10.1038/15446.

Zouboulis CC, Baron JM, Böhm M, Kippenberger S, Kurzen H, Reichrath J, Thielitz A. 2008. Frontiers in sebaceous gland biology and pathology. Exp Dermatol. 17(6):542–551. doi: 10.1111/j.1600-0625.2008.00725.x.

