## Supplemental Materials for "Single-cell analysis of 2,3,7,8-tetrachlorodibenzo-*p*-dioxin (TCDD)-treated murine skin demonstrates that sebaceous gland differentiation precedes *Ahr*-dependent seboatrophy with elevated expression of infundibular *Blimp1*"

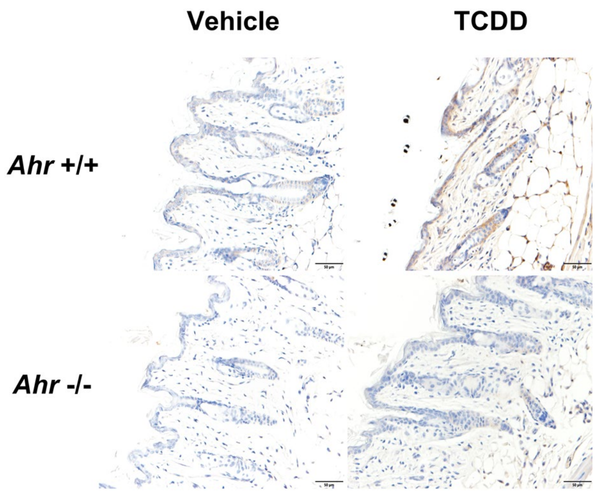


**Figure S1.** Representative images for AHR protein expression at P21**.** Time-mated heterozygous *Ahr* allele dams were treated with either vehicle (corn oil) or TCDD (5 µg/ kg bw) by oral gavage and *Ahr +/+* or *Ahr -/-* pup skin tissue (P21) was stained using an anti-AHR antibody (see materials and methods)**.** Scalebar = 50 µm**.**

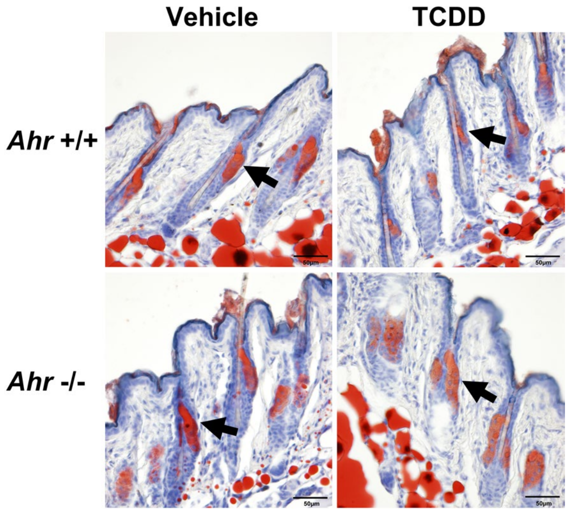


**Figure S2.** Representative images of Oil Red O staining of P21 samples. Time-mated heterozygous *Ahr* allele dams were treated with either vehicle (corn oil) or TCDD (5 μg/ kg bw) by oral gavage and *Ahr* +/+ or *Ahr* -/- pup skin tissue (P21) was stained for lipids with Oil Red O. Arrows point toward the sebaceous glands. Scalebar = 50 μm.

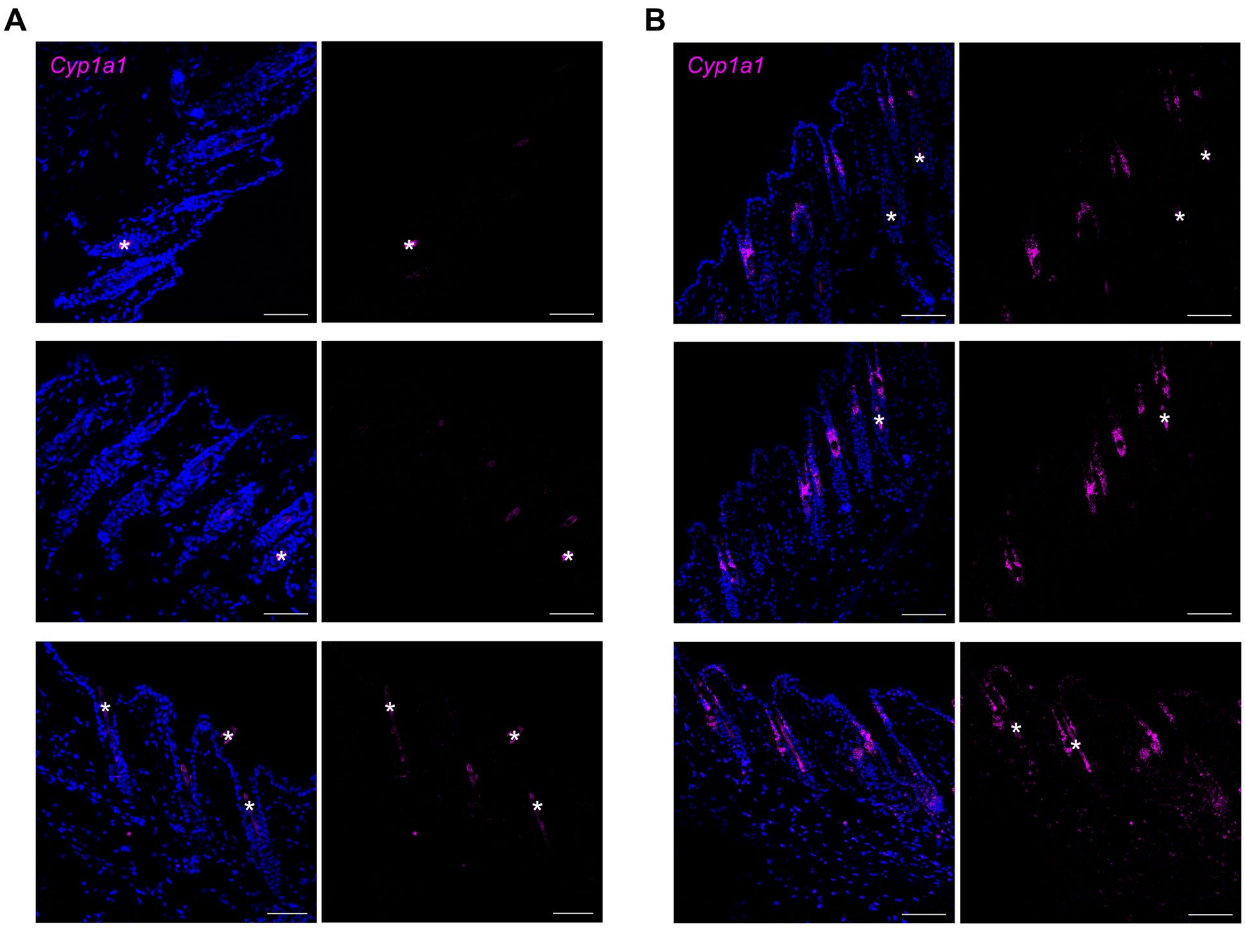


**Figure S3.** *Cyp1a1* expression in skin of *Ahr* +/+ mice treated with **(A)** vehicle (corn oil) or **(B)** TCDD (5 µg/ kg bw); three representative P21 samples per treatment. Scalebar = 50 µm. Hair shaft is indicated by an asterisk.

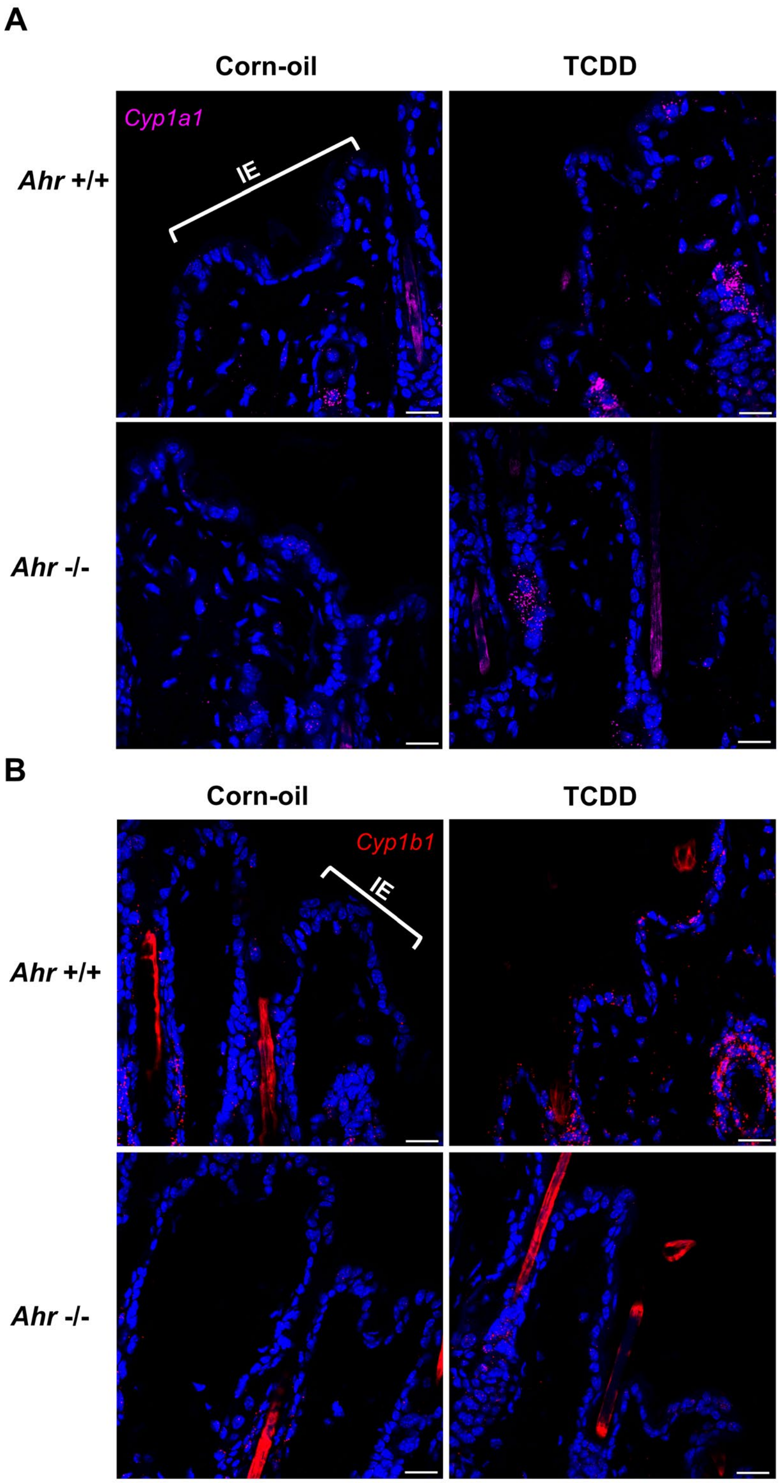


**Figure S4.** *Cyp1a1* and *Cyp1b1* expression in the interfollicular epidermis (IE) at P21. A representative IE is indicated in the first image of each panel. **(A)** *Cyp1a1* expression in the IE is similar across all groups. **(B)** *Cyp1b1* expression in IE. Note: In **Figure 2 (M-N),** *Cyp1b1* expression is significantly higher in the TCDD-treated *Ahr* +/+ samples. Images are a magnified view. Scalebar = 20 μm.

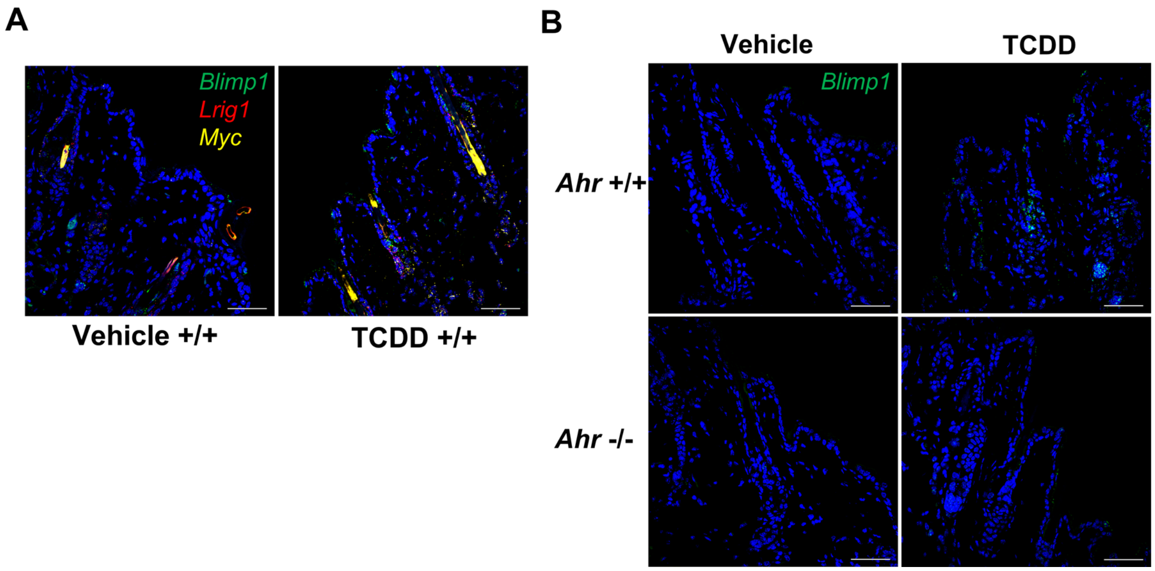


**Figure S5.** Full-sized original images, corresponding to **Figure 3**. **(A)** mRNA Expression of *Blimp1*, *Myc* and *Lrig1* in vehicle- and TCDD-treated skin. **(B)** *Blimp1* expression in the lower infundibulum (IF) at P21. Scalebar = 50 µm**.**

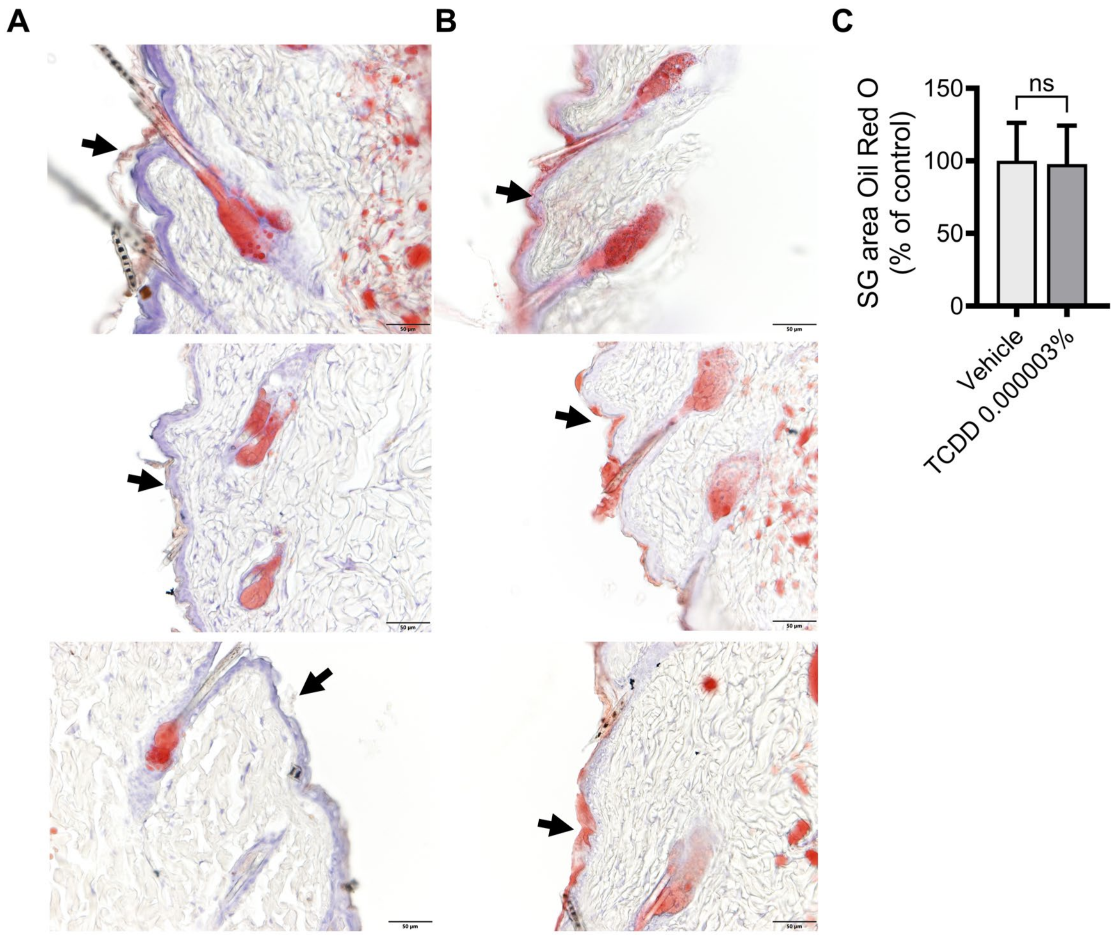


**Figure S6.** Representative Oil Red O staining of frozen skin sections showing increased lipid levels in the interfollicular epidermis (IE). Black arrows point toward the IE. C57BL/6J mice were treated topically once per day for 3 days with either **(A)** vehicle (DMSO:acetone, 1:9) or **(B)** 0.000003% TCDD. Each column represents 3 different samples from the respective treatment group, and each row shows vehicle and treated samples stained at the same time. Scalebar = 50 µm. **(C)** SG area is shown as the percentage of vehicle (100%), mean ± SD (n=3). Statistics were performed using a two-tailed Student's t-test, ns, not significant.

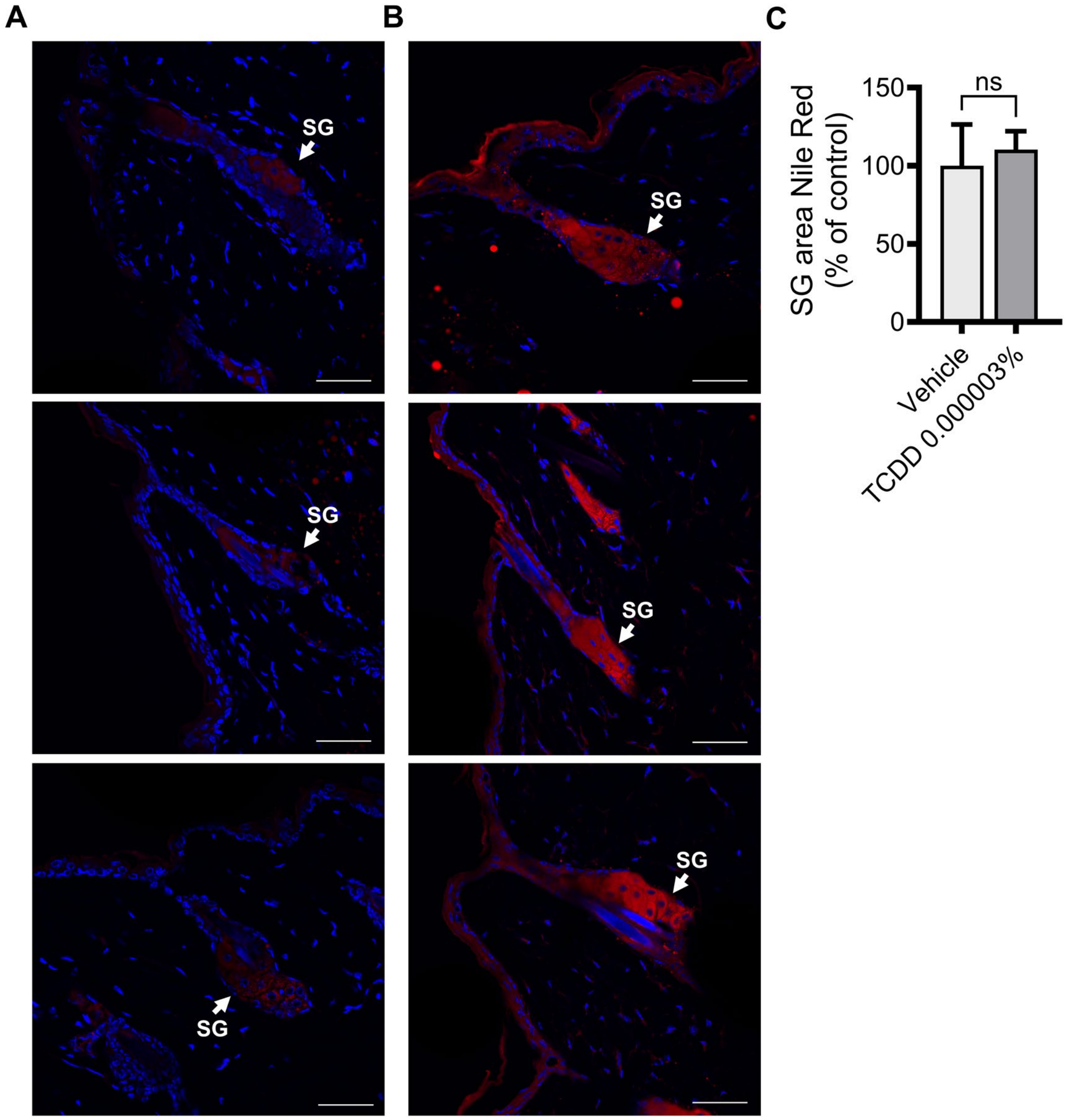


**Figure S7.** Representative Nile Red staining of frozen skin sections showing increased lipid levels in the interfollicular epidermis (IE) and the sebaceous gland (SG). C57BL/6J mice were treated topically once per day for 3 days with either **(A)** vehicle (DMSO:acetone, 1:9) or **(B)** 0.000003% TCDD. Scalebar = 50 µm. **(C)** SG area (mean ± SD) is expressed as the percentage of control (100%). Each symbol represents an individual animal (n = 3). Statistics were performed using a two-tailed Student's t-test, ns, not significant.

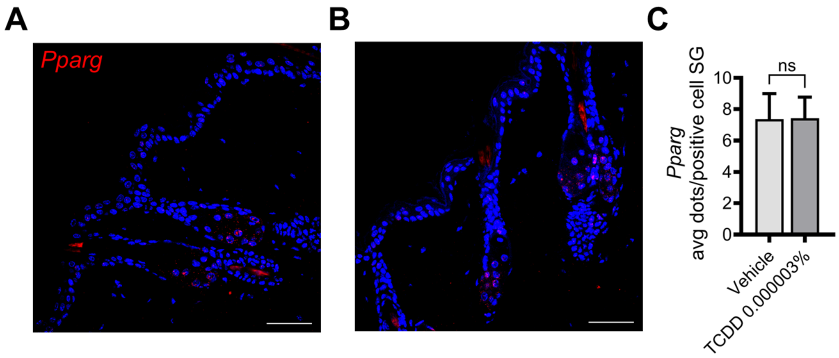


**Figure S8.** *Pparg* expression in a 3-day topical study. C57BL/6J mice were treated topically once per day for 3 days with either **(A)** vehicle (DMSO:acetone, 1:9) or **(B)** TCDD 0.000003% **(C)** Quantification of *Pparg* expressed as average dots per positive cell. The data is shown as mean ± SDs (n = 5). Statistics were performed using a two-tailed Student's t-test, ns, not significant.

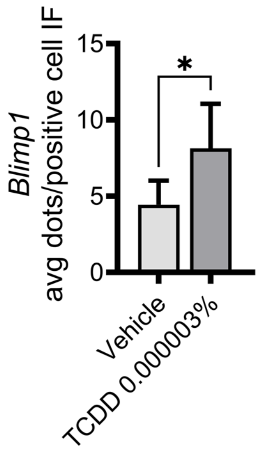


**Figure S9.** Quantification (average dots per positive cell) of *Blimp1* in the lower IF at 3-day. Statistics were performed using a two-tailed Student's t-test, * $p < 0.05$.
